# Dissecting Mechanisms of Ligand Binding and Conformational Changes in the Glutamine-Binding Protein

**DOI:** 10.1101/2023.08.02.551720

**Authors:** Zhongying Han, Sabrina Panhans, Sophie Brameyer, Ecenaz Bilgen, Marija Ram, Anna Herr, Alessandra Narducci, Michael Isselstein, Paul D. Harris, Oliver Brix, Pazit Con, Kirsten Jung, Don C. Lamb, Eitan Lerner, Douglas Griffith, Thomas R. Weikl, Niels Zijlstra, Thorben Cordes

## Abstract

The glutamin-binding protein GlnBP is part of an ATP-binding cassette transporter system in E. coli and uses two well-characterized conformational states, an open ligand-free and a closed-liganded state, to facilitate active amino-acid uptake. Existing literature on its ligand binding mechanism lacked sufficient evidence to univocally assign the kinetic type of binding mechanism for GlnBP: ligand binding prior to conformational change, i.e., an induced fit or the conformational selection, in which the ligand binds the matching conformation from a pre-existing ensemble. Since such mechanistic questions are relevant for our fundamental understanding of how this and other biomacromolecules regulate cellular processes, we here revisit the question for GlnBP. We present a biochemical and biophysical analysis using a combination of calorimetry, single-molecule and surface-plasmon resonance spectroscopy and molecular dynamics simulations. We found that both apo- and holo-GlnBP show no detectable exchange between open and (semi-)closed conformations on timescales between 100 ns and 10 ms and that ligand binding and conformational changes in GlnBP are correlated. A global analysis of our experimental results suggests that the conformational selection model is only compatible with GlnBP for the extreme scenario of very fast conformational exchange between the open and closed states on timescales <100 ns. In contrast all data remains compatible with an induced-fit mechanism, where the ligand binds GlnBP prior to conformational rearrangements. Importantly, our work demonstrates that it is an intricate task to identify the type of kinetic binding mechanism and that this requires not only a sufficient set of data, but also an integrative experimental and theoretical framework to address the question. Based on this concept, we propose that various protein systems, for which so far only insufficient kinetic data are available, should be revisited.

## INTRODUCTION

Periplasmic substrate-binding proteins (SBPs)^1–6^ are small, soluble proteins (molecular weight <100 kDa) that are often associated with membrane complexes, including the superfamily of ATP-binding cassette (ABC) transporters^7^. SBPs recognize and bind numerous classes of substrates including (but not limited to) ions, vitamins, co-factors, sugars, peptides, amino acids, system effectors, and virulence factors^8^. Major biological functions of SBPs are to facilitate membrane transport by delivery of substrate molecules to a transmembrane component or to signal the presence of a ligand^8,9^. They are ubiquitous in archaea, prokaryotes, and eukaryotes and possess a highly-conserved three-dimensional architecture with two rigid domains, D1/D2, that are linked by a flexible hinge composed of β-sheets (Figure 1A), α-helices, or smaller sub-domains^8,9^. The available crystal structures of SBPs reveal that many exist in a minimum of two distinct conformations, a ligand-free open (apo) and a ligand-bound closed state (holo; Figure 1B).

**Figure 1.**
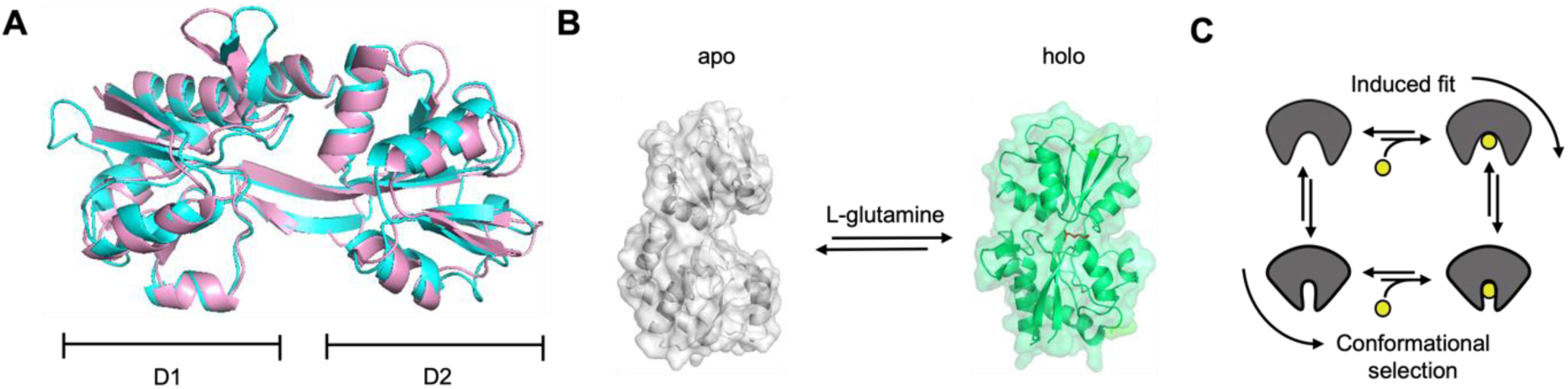
Conformational states and possible ligand binding mechanisms of typical SBPs. (A) Structural comparison of SBD2 from *Lactococcus Lactis* (PDB file:4KR5^10^; cyan) and glutamine-binding protein GlnBP from *E. coli* (pink). SBD2 and GlnBP share 34% sequence identity with a TM-score of 0.90, indicating high structural similarity. (B) Crystal structures of the ligand-free (PDB file:1GGG^11^; grey) and ligand-bound (PDB file:1WDN^12^; green) state of GlnBP from *E. coli*. (C) Sketch of ligand binding via induced-fit (IF) and conformational selection (CS).

Several recent studies focused on the characterization of structural dynamics and conformational heterogeneity as well as the ligand-binding mechanisms that underlie SBP function^13–21^. Based on crystal structures, it was proposed that SBPs use a venus-fly-trap ligand-binding mechanism in which binding traps the ligand via its closed conformation^8,9^. Provided that both ligand binding and conformational changes are well separated in their timescales, the venus-fly-trap corresponds to an induced-fit (IF) ligand binding mechanism (Figure 1C). In IF the event of ligand binding triggers the functionally-relevant conformational change. This intuitive model was challenged by nuclear magnetic resonance (NMR) based paramagnetic relaxation enhancement (PRE) experiments^22^, molecular dynamics (MD) simulations^23,24^ and X-ray crystallography^25–27^ revealing the existence of unliganded closed- or semi-closed states and their dynamic exchange with the respective open (apo) conformation. Similar findings on ligand-independent conformational changes were presented for the maltose-binding protein, MalE^23,24,28^, histidine-binding protein (HisJ)^29^, D-glucose/D-galactose-binding protein (GGBP)^25,27,30,31^, ferric-binding protein (FBP)^32^, choline/acetylcholine substrate-binding protein (ChoX)^26^, the Lysine-, Arginine-, Ornithine-binding (LAO) protein^33^ and glutamine-binding protein (GlnBP)^18,34,35^. For MalE, NMR techniques revealed a low abundance (<10%) semi-closed state that is in rapid dynamic equilibrium with the open apo conformation on the <50 µs timescale. The existence of closed, unliganded state(s) permits an alternative mechanism via conformational selection (CS)^36^, where conformational changes occur intrinsically and prior to ligand binding. In CS the ligand selects the relevant state, e.g., closed, for binding (Figure 1C). IF and CS represent the simplest kinetic schemes to describe the coupling of conformational changes and ligand (un)binding against which available kinetic data can be tested to falsify the types of ligand binding mechanisms.

Ligand-binding mechanisms have also been the focus of several studies of GlnBP^18,19,37^. GlnBP is part of an ABC transporter system in *E. coli* and binds *L*-glutamine with sub-micromolar affinity^38,39^ and arginine with millimolar affinity^40^. It is monomeric and comprised of two globular domains: the large domain (residues 5 – 84, 186 - 224) and the small domain (residues 90 - 180), linked via a flexible hinge (residues 85 – 89, 181 and 189). GlnBP was crystalized in two distinct conformational states: open (apo, ligand-free)^11^ and closed (holo, ligand-bound)^12,38^ (Figure 1B). GlnBP^18,19,37^ has recently been studied by a combination of single-molecule Förster resonance energy transfer (smFRET), NMR residual dipolar coupling (RDC)^18^ experiments, MD simulations^34,35^ and Markov state models (MSMs)^37^. Based on the results, it was proposed that GlnBP undergoes pronounced conformational changes both in the absence^18^ and in the presence^19^ of substrate, involving a total of four to six conformational states. These findings lead to the interpretation that ligand binding in GlnBP could occur by means of a combination of CS and IF^37^ which we found controversial in light of other existing data demonstrating that NMR experiments on GlnBP do not support the idea of intrinsic conformational dynamics in apo-GlnBP^41^. Furthermore, the observation of multiple GlnBP conformers under apo- and holo-conditions via smFRET^18,19,37^ do not align with findings from MD simulations^42^ and smFRET work of substrate binding domains SBD1 and SBD2 (from the amino acid transporter GlnPQ^20,43^), which structurally resemble GlnBP (Figure 1A; SBD2 shows ∼34% sequence identity with GlnBP, TM-score of 0.90). Finally, ligand binding to the fully-closed state of the GlnBP conformation seems rather unlikely, considering the limited accessibility of the binding site, which is also seen for related proteins such as MalE^44^.

Such controversial findings and arguments reveal a central problem in the study of ligand-binding mechanisms, which is the availability of sufficient experimental evidence to distinguish one mechanism from the other. Importantly, both IF and CS imply temporal order of ligand-protein interactions and conformational changes and thus require kinetic data for univocal identification^36,45–49^. The existence of a ligand-free protein conformation, which structurally resembles a ligand-bound form, is necessary^27,50,51^ but by itself not sufficient evidence for a CS mechanism, as ligand binding may not proceed via this conformation at all. Vice versa, the inability to experimentally detect ligand-free closed conformations cannot be taken as an indicator for IF as a dominant pathway^36,45–49^ since only very few techniques are able to detect low abundance (high free energy) conformers and their exchange kinetics with the stable ones. Whether, e.g., a ligand-free closed (or near-closed) conformation^52^ can be observed depends on the magnitude of its equilibrium probability^20^ as well as the sensitivity of the techniques used to probe it. While single-molecule fluorescence approaches can provide such information^14–20^, they often suffer from photon-limited time-resolution and potential labelling artefacts. Analysis of ensemble-averaged relaxation rates from nonequilibrium stopped-flow kinetics or equilibrium NMR experiments can provide the appropriate time resolution, but may be inconclusive under certain experimental conditions^49,53^, e.g., under the pseudo-first-order condition of high ligand concentrations in stopped-flow experiments^36,45–49^. Consequently, to validate or rule-out the presence of a certain ligand-binding mechanism such as IF and CS, a set of complementary and consistent structural, thermodynamic, and kinetic data of the protein system is required in combination with theoretical model building^49^.

Here, we revisit the question of the ligand-binding mechanisms for GlnBP by biochemical and biophysical analyses of ligand binding and its coupling to conformational changes. For this, we used a combination of isothermal titration calorimetry (ITC), smFRET, surface-plasmon resonance (SPR) spectroscopy and MD simulations to derive sufficient evidence that can support the (in)compatibility of the data with any of two kinetic mechanisms. Using smFRET, we observed that apo- and holo-GlnBP show no detectable exchange with other conformational states on timescales between 10 ms and 100 ns. Any observed FRET dynamics could be traced back to photophysical origins rather than to conformational changes. Importantly, in all our smFRET assays, ligand binding and conformational dynamics are highly correlated. In MD simulations of GlnBP, we observed ligand unbinding only after or during the transition from the closed to the open protein conformation. A global analysis of our experimental results suggests that CS model is only compatible with GlnBP for the extreme scenario of very fast conformational exchange between the open and closed states on timescales <100 ns. In contrast all data remains compatible with an induced-fit mechanism, where the ligand binds GlnBP prior to conformational rearrangements.

## RESULTS

### Biochemical characterization of GlnBP and ligand binding

For our study, we produced wild-type protein GlnBP (GlnBP WT) and two double-cysteine variants for analysis of conformational states via smFRET: GlnBP(111C-192C) with point mutations at V111C and G192C (Figure 2—figure supplement 1), and GlnBP(59C-130C) with point mutations at T59C and T130C (the latter was adapted from refs.^18,19^; Figure 2—figure supplement 1C). All protein variants were expressed in *E. coli* and purified using affinity chromatography (see Methods for details). Protein purity was assessed by Coomassie-stained SDS-PAGE analysis (Figure 2A). As reported previously, GlnBP co-purifies with bound glutamine^54^, which was removed by unfolding and refolding of the purified protein. We verified the monomeric state and proper folding of the resulting protein using size-exclusion chromatography (SEC, Figure 2B) by comparing the elution volume and shape of the monodisperse peak of GlnBP before and after the procedure (Figure 2—figure supplement 2).

**Figure 2.**
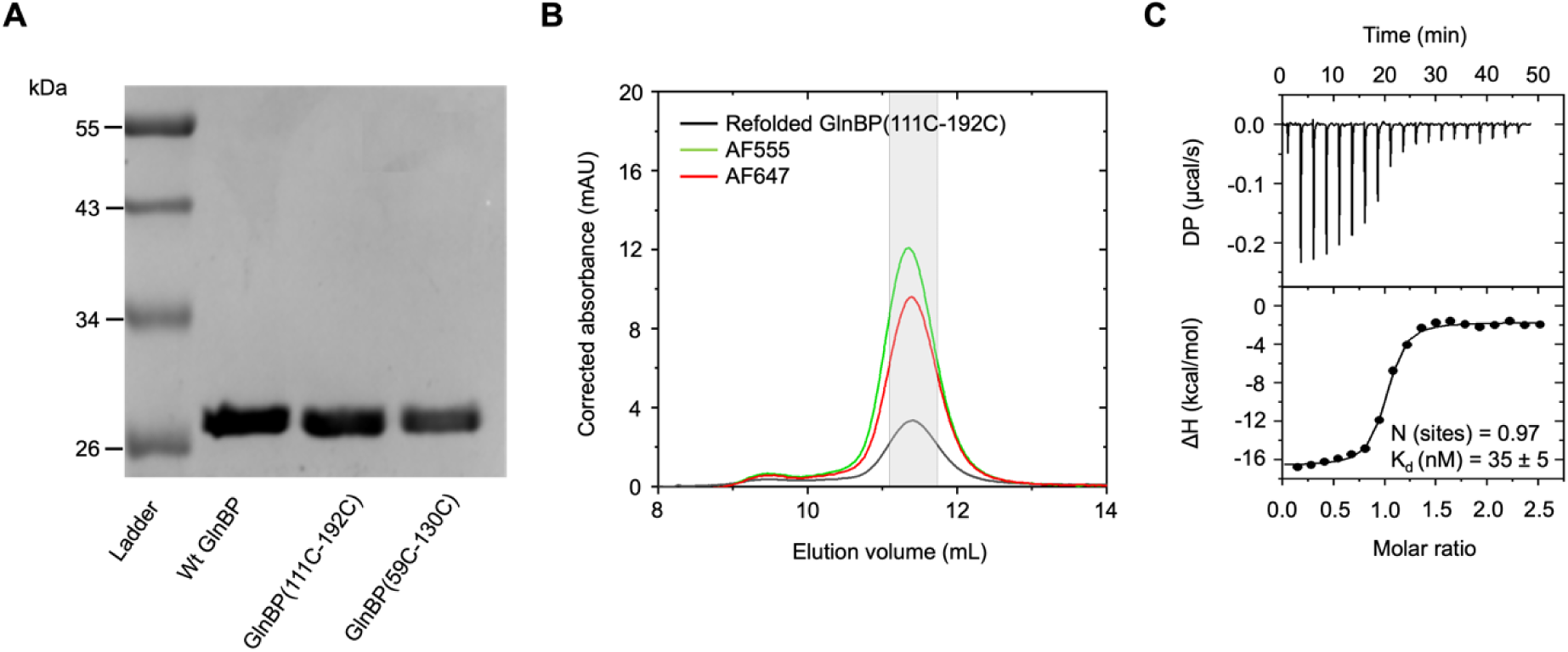
**Biochemical characterization, fluorescence labeling and thermodynamic characterization of ligand binding of GlnBP**. (A) SDS-PAGE analysis of GlnBP purity with coomassie-staining. Lane 1, molecular mass ladder with sizes of proteins indicated in kDa; lane 2, purified GlnBP WT; lane 3, purified double-cysteine variant GlnBP(111C-192C); lane 4, purified double-cysteine variant GlnBP(59C-130C). (B) SEC was used to further purify the fluorescently-labeled proteins. The protein absorption was monitored at 280 nm (black curve), the donor dye absorption (AF555) at 555 nm, and the acceptor dye absorption (AF647) at 647 nm. The labeling efficiency of AF555 and AF647 were estimated to be about 71% and 59%, respectively. For the solution-based smFRET measurements, the used protein fractions are indicated in grey. (C) Ligand-binding affinities of refolded, unlabeled GlnBP(111C-192C) was determined by ITC with a K_d_ = 35 ± 5 nM for L-glutamine (mean value from N = 3 with standard deviation), which is in agreement with previous reports^40^. The free energy of binding was ΔG = - 42.6 kcal/mol with the enthalpy ΔH = −62.3 kcal/mol and entropy contributions - T*ΔS = 19.9 kcal/mol.

To assess the binding affinity of GlnBP WT and the two GlnBP cysteine variants for *L*-glutamine, we performed isothermal titration calorimetry (ITC)^55^. Refolded GlnBP WT showed a K_d_ for L-glutamine of 22 ± 7 nM (Figure 2—figure supplement 3A) and K_d_ values of 31 ± 3 nM and 35 ± 5 nM for the two cysteine variants (Figure 2C, Figure 2—figure supplement 3B). These values are in overall agreement with previously published data^40^. This verifies that the unfolding and refolding process as well as cysteine substitutions did not impact the biochemical properties of unlabeled GlnBP.

### Analysis of conformational states of freely-diffusing GlnBP via smFRET

After assessing the thermodynamic properties of GlnBP, we characterized the conformational states and changes associated to ligand binding via smFRET. With smFRET, it is possible to study biomacromolecules in aqueous solution at ambient temperature, and identify conformational changes, heterogeneity, small sub-populations and determine microscopic rates of conformational changes^56–58^. We performed smFRET experiments on freely-diffusing (Figure 3) and surface-immobilized GlnBP using the refolded variants GlnBP(111C-192C) and GlnBP(59C-130C) labeled with two different dye pair combinations, AF555/AF647 and ATTO 532/ATTO 643, to assess any position- and fluorophore-dependent effects. The smFRET assays were designed such that the inter-dye-distance of the apo state results in a lower FRET efficiency as compared to the holo state of the protein (Figure 2—figure supplement 1A/C).

**Figure 3.**
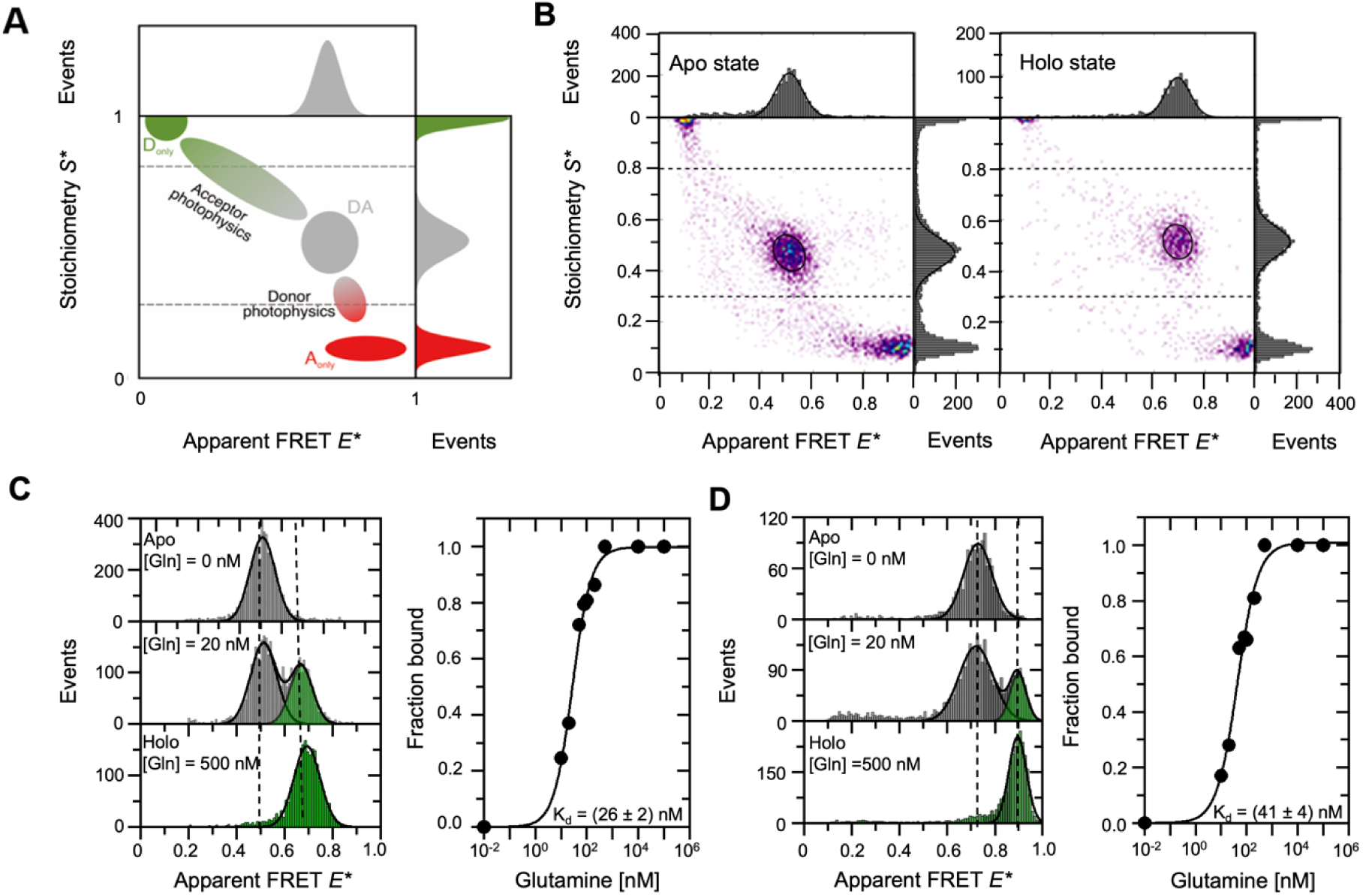
smFRET analysis of GlnBP using diffusion-based µsALEX. (A) Graphical depiction of an E*-S* histogram obtained by μsALEX; panel adapted from ref. 110^60^. Using μsALEX, the stoichiometry S* can be used to separate donor-only (S > 0.8, D_only_), acceptor-only (S < 0.3, A_only_), and the FRET molecular species with both donor and acceptor fluorescently-active fluorophore (S* between 0.3-0.8, DA). Bridge artifacts or smearing caused by donor or acceptor photophysics (photoblinking and/or photobleaching) can cause artificial broadening of the FRET population or a shift of the extracted mean apparent FRET efficiency. (B) μsALEX-based E*-S* histograms of the refolded GlnBP(111C-192C) double-cysteine variant labeled with AF555 and AF647. (C, D) Diffusion-based single-molecule analysis and ligand-binding affinity measurements with μsALEX of doubly-labeled GlnBP(111C-192C) (C) and GlnBP(59C-130C) variants (D) at different ligand concentrations. Values provided are mean +/-SD (N = 3). For plotting purposes the concentration of glutamine in the apo state was set artificially to a value of 0.01 nM in the right parts of panels C/D. Data fitting of the fraction of the high-FRET subpopulation as a function of ligand concentration was performed with the Hill equation, which is a valid approximation for describing the bound fraction of GlnBP as a function of glutamine in the case where [GlnBP] << K_d_.

Solution-based µsALEX^57^ data of GlnBP(111C-192C) labeled with AF555/AF647 are shown in Figure 3B after an all-photon burst search^59^. Both apo and holo states, in the absence and presence of saturation levels of glutamine, respectively, show a clear predominant population of donor-acceptor-labeled protein at S*-values of ∼0.5, with two distinct mean apparent E* values for the apo (mid FRET, 0.51) and holo (high FRET, 0.68) states (Figure 3B/C, Figure 3—figure supplement 1A). This can be interpreted as a transition from open (apo) to closed (holo) GlnBP conformations upon the addition of the ligand. Similar results were obtained for the second double-cysteine variant (GlnBP(59C-130C), Figure 3D, Figure 3— figure supplement 2) and from measurements with a different pair of fluorescent dyes for GlnBP(111C-192C) (ATTO 532/ATTO 643; Figure 3—figure supplement 1B). Notably, further analysis and a comparison of mean accurate FRET efficiencies and the inter-dye distances, show good agreement with simulated inter-dye distances of ±0.2 nm using the open- and closed-GlnBP crystal structures except for the holo-state of GlnBP(59C-130C), which deviated by ∼0.8 nm (Figure 2—figure supplement 1 B/D).

Importantly, a quantitative analysis of the fraction of the closed state (high-FRET) subpopulation as a function of ligand concentration (Figure 3C, D) with a simple binding isotherm and no cooperative effects (n = 1) provides K_d_ values in the 20-50 nM range for all labeled GlnBP variants. These results are fully consistent with ITC (Figure 2C, Figure 3—figure supplement 1/3). Interestingly, we found that arginine, the non-cognate ligand of GlnBP, induces hardly any FRET shifts at even at millimolar concentrations of ligand (Figure 3—figure supplement 4) despite its binding to GlnBP at these concentrations (Figure 3—figure supplement 5).

We were also unable to identify a clear high-FRET subpopulation in the absence of a ligand, which would indicate slow intrinsic exchange of apo/open GlnBP with a (partially) closed conformation on timescales slower than the burst duration, >10 ms (Figure 3B, apo). To estimate an upper bound of the fraction of poorly sampled low abundance states, we determined the percentage of bursts outside of the main FRET population in the range of <E*>±σ of the characteristic FRET population of apo GlnBP(111C-192C). For this, the FRET populations were fitted with Gaussian functions (with mean values and σ), which serves as a good approximation for mean E* values. For representative data sets of AF dyes, we found ∼12% bursts outside of the main peak range (E*_holo_ = 0.64, σ =0.061, N = 626; E*_apo_ = 0.47, σ =0.070, N = 5,013) and for ATTO dyes, ∼4% bursts outside the peak region (E*_holo_ = 0.56, σ =0.056, N = 124; E*_apo_ = 0.37, σ =0.047, N = 2,908). This suggests an upper bound of 5-10% for a subpopulation of other FRET subpopulation, e.g., partially closed conformations of GlnBP. Thus, our results agree with the idea that GlnBP mainly exists in a one state – the open conformation – in the absence of its ligands.

### Screening for rapid conformational dynamics via analysis of “within-burst” FRET dynamics

Next, we analyzed our smFRET data for “within-burst” dynamics using burst-variance analysis (BVA)^61^, multi-parameter photon-by-photon hidden Markov modeling (mpH^2^MM)^62^, intensity-based FRET efficiency versus donor lifetime (E-τ; E stands for FRET efficiency, τ is lifetime) plots^63^ and burst-wise fluorescence correlation spectroscopy (FCS). These analyses provide access to FRET-dynamics that occur on timescales from a few milliseconds down to the sub-µs regime. This allows us to assess whether the observed FRET populations represent stable conformational states or time averages of (rapidly) interconverting states.

We first performed BVA of GlnBP(119-192) data with ATTO 532/ATTO 643 as a dye pair using a dual-channel burst search (DCBS)^59^. In BVA, within-burst E*-dynamics are identified as an elevated standard deviation of the apparent FRET efficiencies, σ(E*), beyond what is expected from photon statistics, i.e., σ(E*) values larger than the theoretical semicircle (Figure 4—figure supplement 1-3, panels A). Our analysis indicates that, for each of the different ligand concentrations, at least some of the recorded single molecules undergo dynamic changes in E* while diffusing through the confocal spot (Figure 4—figure supplement 1). The within-burst dynamics are more prominent for dyes AF555 and AF647 (Figure 4—figure supplement 2) and become most abundant in the variant GlnBP(59C-130C), which was used in previous studies^18,19,37^ (Figure 4—figure supplement 3). It is important to note that dynamic changes in apparent FRET efficiency can have photophysical origins and do not necessarily confirm the presence of conformational dynamics. For example, the apparent dynamic changes in E* might represent within-burst dynamics between FRET-active sub-populations (i.e., S*∼0.5) and FRET-inactive subpopulations (e.g., donor-only, acceptor-only). Therefore, it is essential to quantify the BVA observed dynamics and identify the corresponding E*-S* subpopulations between which the dynamic transitions occur.

**Figure 4.**
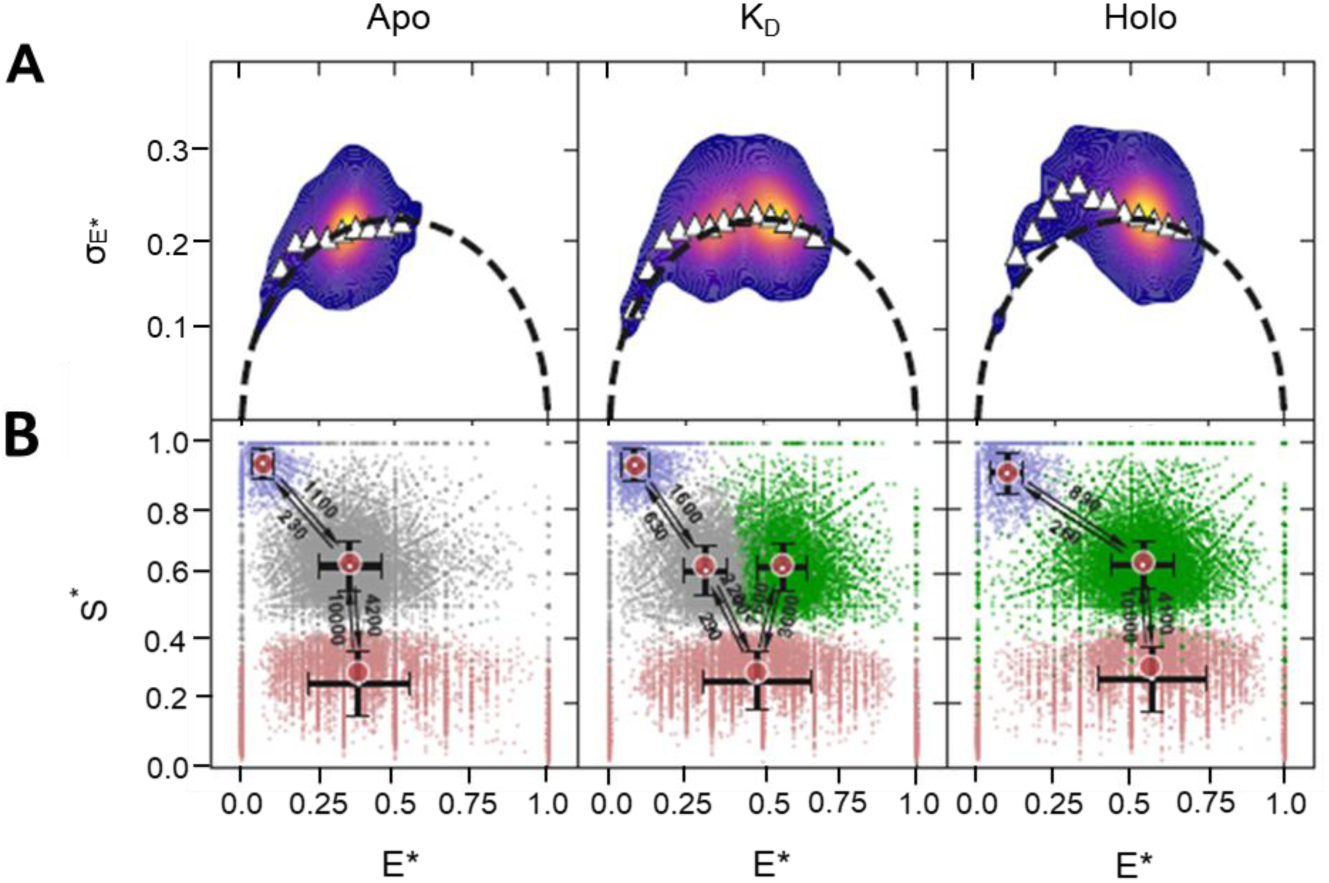
**Screening GlnBP for rapid within-burst FRET dynamics**. (A) Burst Variance Analysis (BVA) showing a weak signature of within-burst FRET dynamics in the low E* regime. (B) Two-dimensional E* versus S* scatter plots of dwells in mpH^2^MM-detected states within bursts detected by the Viterbi algorithm. Arrows and adjacent numbers indicate transition rates in s^-1^. Transitions with rates <100 s^-1^ are omitted since such long dwells in a state before transitions are improbable to occur within single-molecule bursts with durations <10 ms and are most probably a mathematical outcome of the mpH^2^MM optimization framework. The dispersion of the E* and S* values of dwells in mpH^2^MM-detected states are due to the short dwell times in these states, where the shorter the dwell time in a state is, the lower the number of photons it will include, and hence the larger the uncertainty will be in the calculation of E* and S* values of dwells. E* and S* are E* and S* values uncorrected for background, since in mpH^2^MM all burst photons are considered, including ones that might be due to background. Full analysis shown in Figure 4—figure supplement 1.

For this purpose, we used multi-parameter photon-by-photon hidden Markov modelling (mpH^2^MM)^62,64^ to identify the most-likely state model that describes the experimental results based on how E* and S* values change within single-molecule bursts. Such analysis can provide rates of exchange between distinct states of E*/S* and its interpretation is described in detail in Appendix 1. The mpH^2^MM analyses can differentiate whether apparent dynamic changes in E* arise from two conformational sub-populations or from photophysical transitions that do not represent conformational dynamics of GlnBP. Our analysis in Figure 4B shows clear signatures for donor- and acceptor-blinking between bright and dark states of the fluorophores (Figure 4, Figure 4—figure supplement 1-3), i.e., the FRET species with intermediate S* exchange with species of very high and low S* values, respectively. mpH^2^mm identifies single and static apo FRET-active mid-E* state in the absence of a ligand and a single and static FRET-active high-E* state in the presence of saturating levels of ligand, which we ascribe to the open (mid-E*) and closed (high-E*) conformations of GlnBP. It is only in the presence of low concentrations of glutamine (around its K_d_) where two FRET-active sub-populations, representing two distinct conformational states, are identified that might interconvert on timescales slower than 10 ms (i.e., slower than typical burst durations). In conclusion, if intrinsic conformational dynamics existed in apo or holo GlnBP, it could only be between the highly-populated FRET conformation we identify and another conformation that is populated significantly below the sensitivity of our measurement and analysis (i.e., a minor population with a fraction <5-10%) or these transitions would have to occur much faster than the time resolution of our experiments (< 100 µs), which is dictated by the alternation periods in the µsALEX experiment.

To check for the presence of faster dynamics, we used multiparameter fluorescence detection with pulsed interleaved excitation (MFD-PIE)^65^. GlnBP(119-192) labeled with ATTO 532/ATTO 643 were used since this combination showed the least photophysical artifacts. In Figure 5, we first show two dimensional plots of FRET efficiency (E) versus donor fluorescence lifetime values in the presence of acceptor (τ_D(A)_) for apo and holo GlnBP. The theoretical linear relationship between E and τ_D(A)_ defines the static FRET line (Figure 5A, black lines). When the labeled molecules exhibit dynamics faster than the diffusion time, the fluorescence-weighted-average of the donor lifetime becomes biased towards longer donor lifetimes due to the higher brightness values of low-FRET species^63^. Therefore, fast conformational switching is seen as bursts with distinct FRET efficiency values exhibiting a population shift towards the right of the static FRET line. As can be observed from the E-τ plots (Figure 5A), the center-of-mass of the FRET populations for both apo and holo GlnBP are coinciding with the static FRET line, suggesting the absence of conformational changes on timescales faster than milliseconds in line with data in Figure 4.

**Figure 5.**
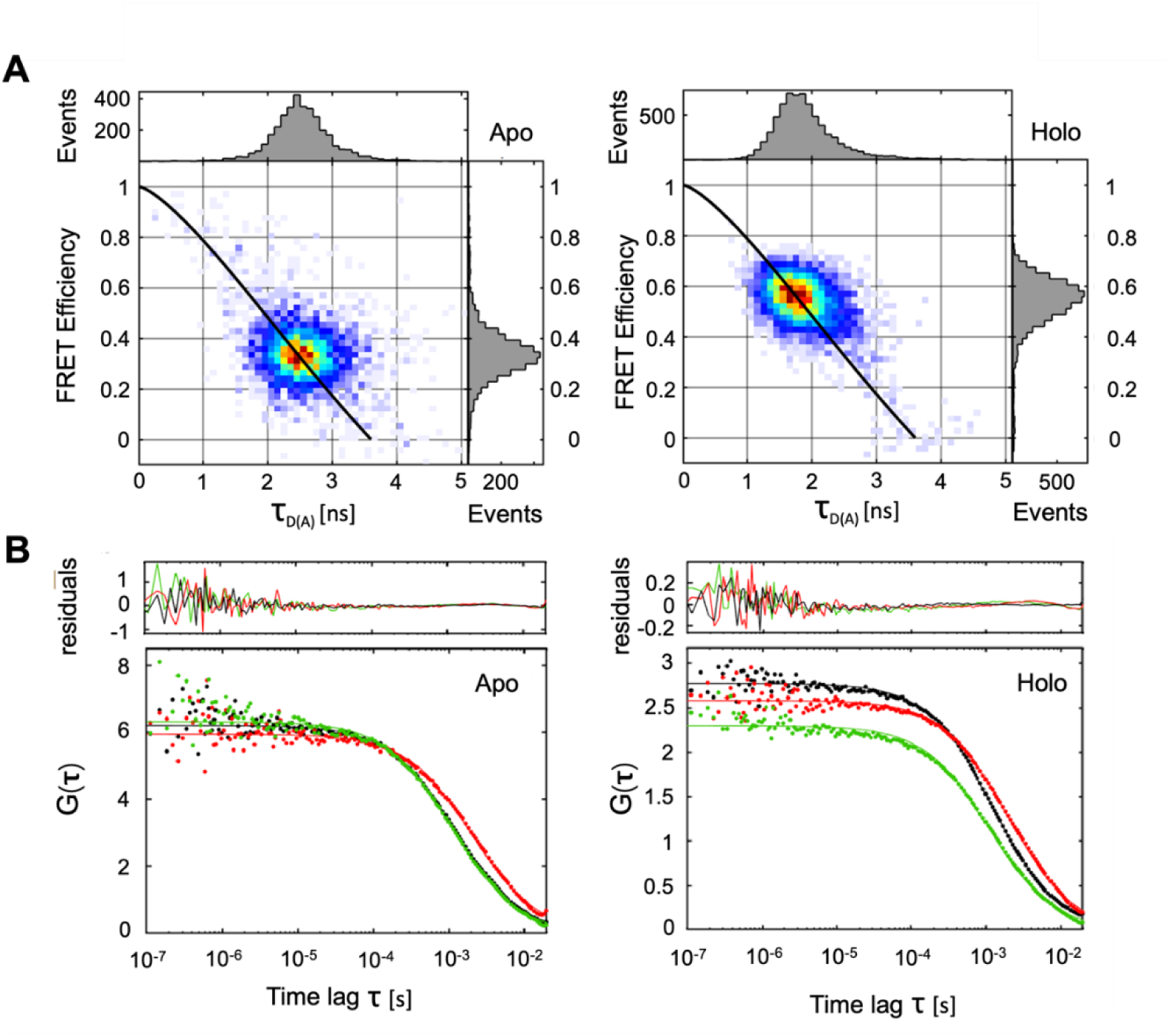
**Screening GlnBP for rapid dynamics within single molecule bursts using E-τ and burst-wise FCS analyses**. (A) Two-dimensional histogram of FRET efficiency (E) versus donor lifetime in the presence of acceptor (τ_D(A)_) for apo (left) and holo (right) GlnBP. The FRET populations coincide well with the theoretical static FRET line (black) indicating the absence of conformational dynamics taking place at timescales faster than ms. (B) Analysis of FRET conformational dynamics using burst-wise FCS for apo and holo states on bursts exhibiting photoactive donor and acceptor fluorophores. The fluorescence autocorrelation functions of the detected donor (DDxDD) and acceptor signal (AAxAA) are displayed in green and red, respectively. The fluorescence cross-correlation function between donor and acceptor signals (DDxDA) is shown in black.

We also looked for dynamics using burst-wise FCS analysis (Figure 5B). For this, bursts containing signal from both fluorophores were selected, padded with 50 ms before and after burst identification and the fluorescence autocorrelation functions of donor (Figure 5B, green curves) and acceptor signals (Figure 5B, red curves) as well as for the fluorescence cross-correlation functions between donor and acceptor signals (Figure 5B, black curves) were calculated. Conformational dynamics are expected to manifest themselves as an anticorrelation contribution in the cross-correlation function between donor and acceptor channels due to fluctuations in FRET efficiencies that occur faster than the translational diffusion component of the correlation functions (∼ 1 ms on our setup)^66^. The burst-wise FCS analysis at times <100 µs resulted in plateaued cross-correlation functions (Figure 5B, black lines) for apo and holo states indicating the lack of dynamics down to the time-resolution of the experiments, i.e., the typical clock time of the photon time tagging on the order of 100 ns. It has to be mentioned that minor population exchange concerning <10% of molecules cannot be excluded with absolute certainty, particularly for the time regime <10 µs. Here, the noise increases due to limited photon budget, yet no clear indication for a cross correlation related to conformational changes are seen, also supported by non-systematic fluctuations in the residuals (Figure 5).

### Studies of surface-immobilized GlnBP via TIRF microscopy

Next, we characterize GlnBP and its conformational dynamics on timescales beyond the residence time of molecules in the confocal excitation volume (i.e., >1-10 ms) with the hope to obtain information on rare conformational events. We consequently conducted smFRET with NTA-based surface-immobilization of the GlnBP His-tag using TIRF microscopy (see Appendix 2 and accompanying Appendix 2 Figures 1-5 for details). We reasoned that this would also allow the direct comparison of our results to those of Wang, Yan and co-workers^18,19,37^. Importantly, in our analysis, we found that various buffer additives used for oxygen depletion have the same effect on GlnBP as the addition of glutamine (i.e., apo-GlnBP becomes artificially “closed” in the presence of the additives) as we demonstrated in solution-based µsALEX experiments (Appendix 2 Figure 1). Consequently, these additives were omitted since their effects mimic that of substrate binding. Strikingly, the conformational states of GlnBP were also partially altered upon surface immobilization (Appendix 2 Figure 2), i.e., the E* values of GlnBP in apo/holo-state were significantly higher than in solution (Appendix 2 Figure 2, 4). Furthermore, GlnBP did not retain its full biochemical activity on the glass coverslips (Appendix 2 Figure 2), i.e., only ∼50 % of all GlnBP molecules showed the expected shift towards higher FRET values upon addition of the ligand (Appendix 2 Figure 2F). To validate our setup and immobilization approach, we additionally tested dsDNA (Appendix 2 Figure 2A, C) and the two previously studied proteins SBD1 and SBD2 (Appendix 2 Figure 5). Here, we did not observe discrepancies in FRET efficiency or biochemical activity, and the data of freely-diffusing and surface-immobilized species were consistent.

Our combined smFRET analysis of GlnBP under different biochemical conditions suggests that conformational changes are tightly coupled to the ligand glutamine (Figure 3). We can also rule out prominent conformational dynamics on timescales between 100 ns and 10 ms of apo and holo GlnBP via mpH^2^MM, MFD-PIE, and burst-wise FCS (Figure 4, 5). Furthermore, our analysis suggests that apo GlnBP does not adopt (partially) closed conformations on the timescale >10 ms with an abundance >5-10 % (Figure 3-5). While these results provide valuable information on ligand binding affinity, conformational heterogeneity and timescales of conformational dynamics in GlnBP, they are insufficient to exclude one or the other kinetic ligand-binding mechanism (IF vs. CS). We thus decided to integrate the obtained information into a general theoretical framework for analysis of ligand-binding mechanisms^36,45,46^, for which additional knowledge of the association and dissociation rates of ligand binding are required.

### Insights on ligand binding kinetics from bulk spectroscopy

Such kinetic information is available from surface plasmon resonance spectroscopy (SPR) or stopped-flow experiments. Since SPR was available to us, we immobilized GlnBP via its His-tag on a sensor chip and monitored its interaction with glutamine as a function of time. Even though GlnBP became partially inactive during immobilization for smFRET in TIRF microscopy (Appendix 2 Figures 1-5), we reasoned that non-functional GlnBP will not be observed in SPR since only functional protein can contribute to the signal changes. The assumption that GlnBP remains functional on SPR-chips was validated by the match of ligand-binding characteristics obtained from ITC (Figure 2C, Figure 2—figure supplement 3), smFRET (Figure 3C, D) and SPR (Figure 6A).

**Figure 6.**
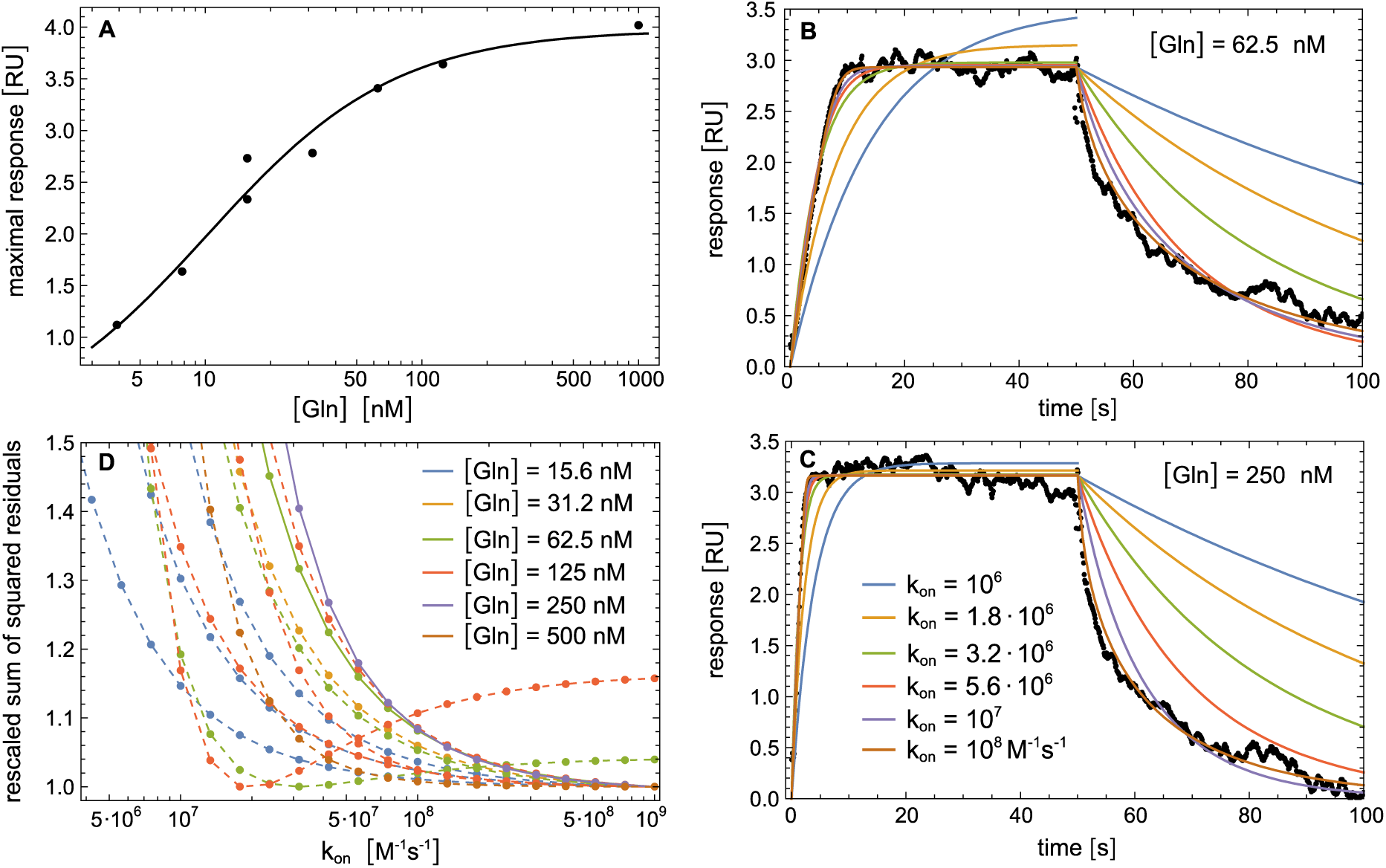
Kinetic analysis of ligand binding and dissociation in GlnBP using SPR. (A) Fitting of maximal responses in sensorgrams from a measurement set with [Gln] concentrations from 7.8 to 1000 nM (data points) with f = c⁄(1 + *K*_*d*_⁄[Gln]) leads to *K_d_* = 10±1 nM. (B, C) SPR sensorgrams with an association phase of 50 s at the indicated glutamine concentrations [Gln] followed by a dissociation phase of 50 s with [Gln] = 0 in the bulk flow (data points), and fits of the sensorgrams with the reaction scheme (1) for different values of the effective on-rate constant *k*_on_ (see Methods for details). (D) Rescaled sum of squared residuals versus *k*_on_ for fits of sensorgrams with different values of [Gln] in the association phase. Note that multiple repeats for the ligand concentrations [Gln] = 15.6 nM, 62.5 nM, and 125 nM are plotted. The two curves with full lines correspond to fits in panels B and C. The 11 curves with dashed lines correspond to the fits in Figure 6—figure supplement 1. Note that panels are arranged in clock-wise order.

In SPR, GlnBP showed specific and stable interaction with glutamine based on the magnitude of the equilibrium RU response as a function of glutamine concentration (Figure 6A). Analysis of the concentration-dependent maximal RU units yields a K_d_ of 10 nM (Figure 6A). The overall maximal response of around 3-4 RU indicates a 1:1 stoichiometry of glutamine assuming a monomeric state of GlnBP (Figure 2C, Figure 6). Kinetic association and dissociation experiments were conducted under pseudo-first order conditions, i.e., the assumption of constant glutamine concentrations during an SPR run, due to the applied flow of buffer. The data were analyzed with the standard two-step reaction scheme^67,68^:

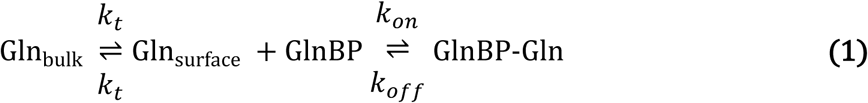

This includes a mass-transport step between the bulk solution of the applied flow and the sensor surface with transport rate *k_t_* in both directions, and a binding step with effective on- and off-rate constants, *k*_on_ and *k*_off_. Because of the dominance of mass transport, fits of this reaction scheme to the SPR sensorgrams using fit parameters *k_t_* and *k*_on_ (after substituting *k*_off_ with *K_d_ k*_on_ in the scheme) do not allow determination of *k*_on_ within reasonable error bounds. However, fits with fixed values of *k*_on_ indicate that effective on-rate constants smaller than 10^7^ M^-1^s^-1^ are incompatible with the sensorgrams (Figures 6B, C and Figure 6—figure supplement 1). More precisely, plots of the rescaled sum of squared residuals for these fits versus *k*_on_ (Figure 6D) indicate a lower bound of at least 3·10^7^ M^-1^s^-1^ for *k*_on_; this implies *k*_off_ = *K_d_ k*_on_ > 0.3 s^-1^ (with K_d_ = 10 nM). Among the 13 plots in Figure 6D, and among the 4 plots for [Gln] = 125 nM, only one plot exhibits a minimum of the sum of squared residuals below this bound and is therefore likely an outlier.

### Sequences of events along MD simulation trajectories

To further investigate the coupling between conformational changes of GlnBP and ligand binding/unbinding, we performed atomistic simulations starting from the ligand-bound GlnBP structure with the AMBER20 software implementation for graphics processing units (GPUs)^69,70^ and the ff14SB force field parameters^67,71^ (see Methods for details). To observe unbinding events on the microsecond timescale accessible in our simulations, we reduced all interactions between the protein and the ligand by 16%. With these reduced interactions, we observed ligand unbinding and a conformational change from the closed to the open GlnBP conformation in 5 out of 20 simulation trajectories with a length of 2 µs. Figure 7 illustrates characteristic distances for GlnBP opening and ligand unbinding on these 5 trajectories for time windows of 500 ns around the unbinding point. GlnBP opening is monitored by the distances between the C-α atoms of the residues 117 and 137 in domain 2 and the residue 51 in domain. We chose these residue pairs because they exhibit large relative changes in distance, with distances of 4.5 and 7.5 Å between residues 51 and 117 and residue 51 and 137 in the closed GlnBP conformation, respectively, and distances between about 15 and 30 Å in the open conformation for both pairs. Ligand unbinding is monitored by the root mean square deviation (RMSD) between the non-hydrogen atoms of the ligand in the simulation structures and the ligand in the bound crystal structure, after alignment of either the D1 or the D2 protein domain of these structures. These two RMSDs quantify the distance of the ligand to its native binding position on D1 and D2, respectively.

**Figure 7.**
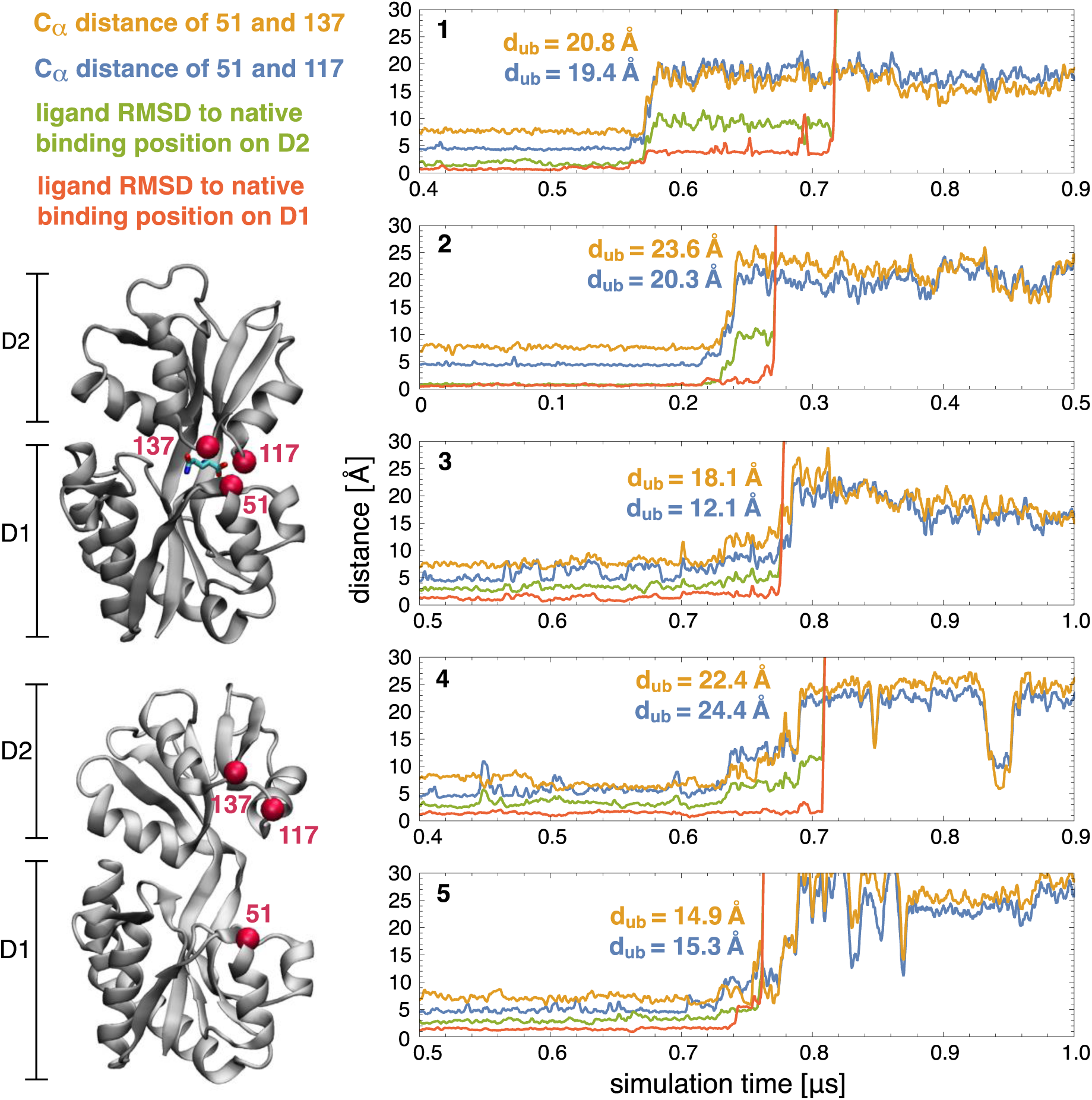
Protein conformational changes and ligand unbinding along MD simulation trajectories. Characteristic distances reflecting protein opening (blue, yellow) and ligand unbinding (green, red) within time windows of 500 ns around the unbinding point of 5 out of 20 trajectories with a total length of 2 µs starting from the closed protein-ligand complex. On the 15 other trajectories, the protein remained in the closed ligand-bound state. To observe unbinding on the microsecond timescales accessible in the simulations, the interactions between the protein and ligand were reduced by 16% in the simulation model (see Methods). The distances d_ub_ are the distances between the C_α_ atoms of the residues 51 and 117 (blue) and 51 and 137 (yellow) at the ligand unbinding point, i.e. at the time point at which the ligand RMSD to the native binding position on domain D1 (red) exceeds 10 Å.

In trajectories 1, 2, and 4 in Figure 8, ligand unbinding occurs clearly after the opening transition of the protein, in agreement with the induced-fit pathway of Figure 1C. During the opening transition of these trajectories, the ligand RSMD to the native binding position on D2 increases, which reflects the breaking of the ligand contacts to D2 during opening. The ligand RSMD to the native binding position on D1 remains low until the unbinding point, at which also the ligand contacts to D1 break. On the trajectories 3 and 5, in contrast, the ligand already unbinds during the opening transition of the protein, but also only after substantial opening at distances d_ub_ of the residues 117 and 137 to residue 51 at the unbinding point that are much larger than the corresponding distances in the bound conformation. It is important to note that the reduction of the protein-ligand interactions in our simulations lowers the binding free energies of the two protein conformations rather homogeneously, akin to a reduction of the ligand concentration, and reducing the ligand concentration is known to shift the flux towards the conformational-selection route of Figure 1C^72–74^, if parallel pathways are possible.

**Figure 8.**
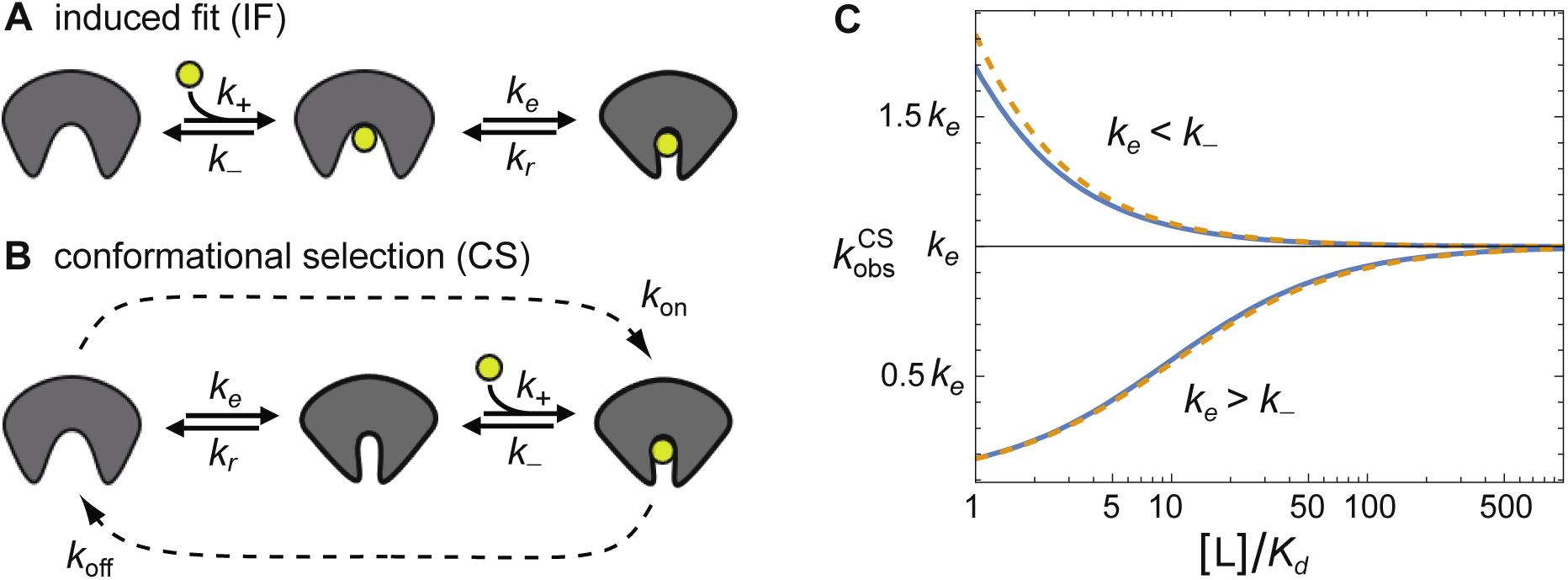
Kinetic description of IF and CS pathways and dominant relaxation rate 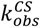 ***of the CS pathway*_s_.** (A, B) Induced-fit and conformational-selection pathways with conformational excitation and relaxation rates, *k_e_* and *k_r_*, and with association and dissociation rate constants, *k*_+_ and *k*_-_, for the binding-competent conformation of the pathway. (C) Dominant relaxation rate, 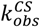, of the conformational-selection pathways versus ligand concentration [L]. Blue lines represent the exact pseudo-first-order result 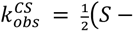 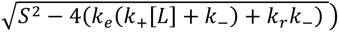 with *S* = *k*_*e*_ + *k*_*r*_ + *k*_+_[*L*] + *k*_−_ and *K*_*d*_ = *k*_−_(*k*_*e*_ + *k*_*r*_)⁄*k*_+_*k*_*e*_ for *k*_−_ = 10 *k*_*e*_ and *k*_*r*_ = 9 *k*_*e*_ (upper curve) and *k*_−_ = 0.1 *k*_*e*_ and *k*_*r*_ = 9 *k*_*e*_ (lower curve). The dashed yellow lines represent the approximate result from equation **2**. For the induced-fit pathway, the dominant relaxation rate 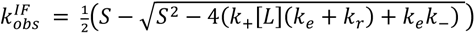 with S as above is monotonously increasing (similar to 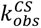 for *k*_*e*_ > *k*_−_) and has the limiting value *k*_*e*_ + *k*_*r*_ at large ligand concentration^36,46^.

Based on the simulation IF seems the dominant binding mechanism also for the original, un-rescaled protein-ligand interactions of our atomistic model, because the sequences of opening and unbinding events observed on our simulations at weakened interactions clearly point towards induced fit, and because the weakening of the interactions rather decreases than increases the tendency for induced fit in Figure 1C. For the original, un-rescaled protein interactions, we expect significantly longer dwell-times in the closed state, significantly longer times for ligand unbinding from D1 after domain unbinding compared to the trajectories 1, 2, and 4 of Figure 7, and a significantly lower probability for ligand unbinding already at protein unbinding as on trajectories 3 and 5.

In addition, we performed simulations starting from the closed GlnBP structure with removed ligand to explore the conformational dwell times in the exchange between the closed and open conformation in the ligand-free state. We observed transitions from the closed to the open GlnBP conformation on 11 out of 20 MD simulation trajectories with a length up to 3 µs (see Figure 7 – figure supplement 1). On the remaining 9 trajectories, the closed conformation persisted for the simulation length of 3 µs. The fraction P(t) of trajectories that exhibit an opening transition up to timepoint t points towards a mean dwell time of several hundred nanoseconds for the closed conformation in the ligand-free state.^75^ On the 11 trajectories that exhibited opening transitions, no subsequent transitions back to closed conformation were observed, which indicates clearly longer dwell-times in the open ligand-free GlnBP conformation of the simulations.

## DISCUSSION & CONCLUSION

Conformational states of macromolecular complexes and changes thereof govern numerous cellular processes including replication^76^, transcription^77,78^, translation^79^, signal transduction^80–82^, membrane transport^83,84^, regulation of enzymatic activity^85–88^, and the mode of action of molecular motors^89,90^. While many conformational changes that are triggered by ligand binding have been characterized extensively, it has also become evident that proteins exhibit prominent intrinsic structural dynamics without the involvement of ligands or other biomacromolecules^1–6,91–96^. Elucidating the kinetic binding mechanisms of proteins and biomolecules will advance our understanding of their fundamental mechanisms and allow the identification of critical steps that might allow rational design of selective and effective inhibitors.

In a four-state system (Figure 1C), ligand-binding can occur via two ‘extreme’ kinetic pathways, i.e., ligand binding occurs before conformational change (induced fit, IF) or conformational change occurs before ligand binding (conformational selection, CS). The clear temporal ordering of ligand binding and conformational change along either of these pathways implies that the binding transition time, i.e., the time the ligand needs to enter and exit the protein binding pocket, are small compared to the dwell times of the protein in the two conformations. An important notion is that the ligand binding mechanisms IF/CS only require temporal separation of ligand binding and conformational changes and are independent of the type of conformational motion found in the specific protein. While the concrete conformational motion can be distinct for different SBPs, e.g., a one- or two-domain motion, spring hammer type of motion^115^, the type of conformational motions need not be confused with a kinetic ligand-binding mechanism IF/CS.

For GlnBP we dissected the ligand-binding processes and conformational dynamics using complementary techniques. We used smFRET experiments to monitor dynamics of conformational changes, SPR to monitor ligand binding and dissociation kinetics, we obtained ligand affinity values from ITC, SPR and smFRET, and explored sequences of conformational opening and ligand unbinding events on simulation trajectories starting from the bound complex. GlnBP fulfils all criteria to use either an IF or CS mechanism since the essential temporal ordering of binding and conformational changes is plausible for SBPs due to their small ligands^45^ and confirmed by the simulation data in Figure 7. Since IF/CS represent the simplest kinetic schemes to describe the coupling of conformational changes and ligand (un)binding, we firmly believe that testing available data against these should be the first step before constructing more complex networks of states.

We consequently ask the question, which binding mechanism is compatible with all the data. We hereby follow a published theoretical framework that aims at an unambiguous assignment of the reaction schemes via kinetic rate analysis^36,45,46^. In essence, we test whether the experimental parameters are compatible both with the IF pathway and the CS pathway (Figure 1) or only one of them. Both pathways are shown in Figure 8A, B with the relevant kinetic parameters, i.e., conformational excitation and relaxation rates, *k_e_* and *k_r_*, and with association and dissociation rate constants, *k*_+_ and *k*_-_, for the binding-competent conformation.

Our smFRET analysis indicates that ligand binding is correlated to a conformational change from an open to a closed state of GlnBP and gives detailed information on the conformational dynamics. It excludes prominent structural dynamics of apo- and holo-GlnBP on timescales above 100 ns. While we cannot explicitly rule out conformational exchange of minor sub-populations of potential ligand-free (partially-)closed conformations and apo-GlnBP, we estimate an upper bound for such processes of <10%. The analysis of SPR sensorgrams lead to the bounds *k*_on_ > 3·10^7^ M^-1^s^-1^ and *k*_off_ = *K_d_ k*_on_ > 0.3 s^-1^ (with K_d_ = 10 nM) for the effective on- and off-rate constants *k*_on_ and *k*_off_ at all considered ligand concentrations of glutamine up to 500 nM.

Based on these information, we first discuss the scenario of a dominant CS pathway in GlnBP. To relate it to the effective on- and off-rates of the SRP analysis, we note that the relaxation rate 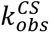 of the CS reaction scheme in Figure 8B can be well-approximated by:

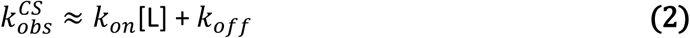

Here, the effective on- and off-rate constants are *k*_𝑜𝑛_ = *k*_*e*_*k*_+_⁄(*k*_*r*_+*k*_+_[L]) and *k*_𝑜𝑓𝑓_ = *k*_*r*_*k*_−_⁄(*k*_*r*_+*k*_+_[L]) that depend on the conformational transition rates, *k_e_* and *k_r_,* between the open and closed conformation in an unbound GlnPB and on the rates, *k*_+_ and *k*_-_, for the binding step in the closed conformation along this pathway. This approximation holds for small populations of the closed conformation in ligand-free GlnPB with upper bound of 10% from the smFRET analysis and for ligand concentrations [L] > *K_d_* and, thus, for all the concentrations shown in Figure 6^45^. At the largest ligand concentration of 500 nM of the SPR sensorgrams, we obtain 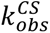 > 15 s^-1^ from this equation, using a lower limit of 3·10^7^ M^-1^s^-1^ for *k*_𝑜𝑛_ . Eqn. 2 can be further simplified to

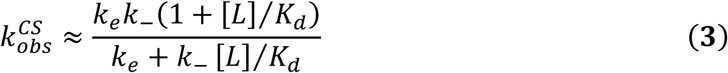

with *K_d_* = *k_-_k_r_*/ *k_+_k_e_*. The limiting value of 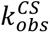 at large ligand concentration [*L*] obtained from this equation is *k_e_*. To conclude the argument, we now consider two cases: (1) for *k_e_ > k_-_*, the relaxation rate 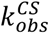 increases with [L] as seen in Figure 8C (lower curve). The limiting value *k_e_* of 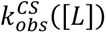 is therefore larger than 15 s^-1^, because 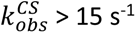 at [*L*] = 500 nM (see above). (2) for *k_e_ < k_-_*, the relaxation rate 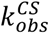 decreases with [L] as seen in Figure 8C (upper curve). In this case, 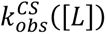 is already very close to its limiting value *k_e_* at [*L*] = 500 nM for *K_d_* = 10 nM. In both cases, we thus obtain *k_e_ >* 15 s^-1^, and from this, *k_r_ > 9 k_e_ >* 135 s^-1^ for an upper bound of 10% of the population *k*_*e*_ ⁄*k*_*e*_ + *k*_*r*_ of the closed conformation in ligand-free GlnBP. However, rates *k_r_ >* 135 s^-1^ correspond to transition timescales <7.4 ms, which are timescales for conformational dynamics of apo GlnBP that are precluded by the smFRET results presented here. Alternatively, timescales smaller than 100 ns are “allowed” for the conformational exchange between the open and closed state, which is theoretically possible considering our MD results (Figure 7).

In contrast to the limited validity of the CS mechanism for very fast exchange between the open and closed state, IF is fully compatible with all experimental data presented here. Similar as for eqn. 2 for IF, we can approximate the relaxation rate *k*^𝐼𝐹^ in SPR:

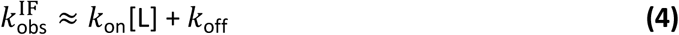

Here the effective on- and off-rate constants are *k*_on_ = *k*_+_*k*_*r*_⁄(*k*_−_+*k*_*r*_) and *k*_off_ = *k*_−_*k*_*e*_⁄(*k*_−_+*k*_*r*_). From the equation for the effective off-rate constant *k*_off_, we obtain which implies

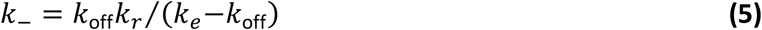

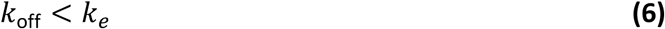

From our SPR results in Fig. 6, we concluded a lower bound of 3·10^7^ M^-1^s^-1^ for *k*_on_ in a range of ligand concentrations [L] from 15.6 to 500 nM, which likely holds also for smaller [L]. Based on stopped-flow mixing experiments of GlnBP and Gln more than five decades ago^9^, an effective on-rate constant of about 10^8^ M^-1^s^-1^ has been obtained from numerical fits of stopped-flow relaxation curves at concentration ratios of 1:1 and 2:1 of Gln and GlnBP. For a plausible range 3·10^7^ M^-1^s^-1^ < *k*_on_ < 10^8^ M^-1^s^-1^ of on-rate constants and *K_d_* values of 10 – 20 nM from different methods (ITC, smFRET, SPR), we obtain 0.3 s^-1^< *k*_off_ < 2 s^-1^ as range for the effective off-rate constant *k*_off_ = *K_d_ k*_on._ Together with Eq. (5), our smFRET results with lower bounds of 100 s^-1^ for the conformational exchange rates *k_e_* and *k_r_* (corresponding to timescales >10 ms) and an upper bound of about 10% for the relative probability 𝑃_OL_ = *k*_*e*_ ⁄(*k*_*e*_ + *k*_*r*_) of conformation OL among the two bound states of GlnBP lead to

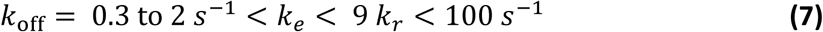

This equation shows that the IF pathway is compatible with our results. Eq. (7) in turn results in a lower bound for 𝑃_OL_of about 0.3 to 2%, and together with Eq. (5), in the lower bounds *k*_−_ ≈ 10 *k*_off_ for 𝑃_OL_ = 10%, *k*_−_ ≈ 20 *k*_off_ for 𝑃_OL_ = 5%, and *k*_−_ ≈ 100 *k*_off_ for 𝑃_OL_ = 2%. Corresponding lower bounds for the on-rate constant *k*_+_ of the binding-competent open conformation of the IF pathway then follow from *k*_+_ = (*k*_−_⁄*K*_*d*_) *k*_*e*_ ⁄(*k*_*e*_ + *k*_*r*_) = (*k*_−_⁄*K*_*d*_)𝑃_𝑂*L*._

We thus consider IF to be the simplest model that correctly describes the ligand binding mechanism in GlnBP in light of the data and simulations presented here, but clearly state that CS remains possible in case that exchange of between the open and closed state in GlnBP is very fast. Another argument to support IF is the notion that the open conformation is more likely to bind substrate than the closed one based on steric arguments (see Appendix 3). A potential improvement in our argumentation would be to include relaxation kinetics^97^ without the mass transport limitations in SPR, which is particularly relevant for small ligand molecules. Thus, stopped-flow (FRET) experiments, which have already been used in the 1970s for binding-rate determination in GlnBP^98^, would be a more direct approach that could complement smFRET data and might lead to more robust as presented above.

What implications do our results and the proposed integrative strategy for determining (or excluding) ligand binding mechanisms have for other protein systems? Generally, we encourage the use of similar strategies for other biomacromolecular systems, and revisiting various SBP systems (and their binding mechanisms). This is relevant since there are many findings and controversial interpretations whenever intrinsic conformational motion or closed-unliganded conformations were identified for the maltose binding protein MalE^23,24,28^, histidine binding protein (HisJ)^29^, D-glucose/D-galactose-binding protein (GGBP)^25,27,30,31^, ferric-binding protein (FBP)^32^, choline/acetylcholine substrate binding protein (ChoX)^26^ and the Lysine-, Arginine-, Ornithine-binding (LAO) protein^33^. Also the advent of single-molecule approaches, such as nanopore-recordings^13^ and single-molecule Förster-resonance energy transfer (smFRET)^14–21^ provided a large pool of data for various ABC transporter-related SBPs^20,43^ with a wide range of distinct ligands such as metal ions^20,100^, osmolytes^20,101^, amino acids^16–21^, peptides^20^, sugars^7,20,62,102,103^, siderophores^104^, and other small molecules^105^ – for most of which additional kinetic data is required to univocally assign a kinetic ligand-binding mechanism.

While SBPs exhibit somewhat conserved structure, certain members show collective differences in structural key features. For example, type I and type II SBPs differ in their overall core topology and in the composition of their hinge domain with two ß-strands for type II, but three strands for the type I family.^99^. Applying our strategy in a comparative study could help to reveal how such hinge-domain differences contribute to conformational dynamics, thereby strengthening the link between proteins secondary structure elements, three-dimensional architecture, and function.

## ACKNOWLEGDEMENTS

This work was financed by the European Comission (ERC-STG 638536 – SM-IMPORT to T.C.), Deutsche Forschungsgemeinschaft (GRK2062, project C03 to T.C.; SFB863, project A13 to T.C. and Sachbeihilfe CO 879/4-1 to T.C., Project 449926427 to K.J., SFB1035 (201302640, project A11 to D.C.L.), the Bundesministerium für Bildung und Forschung (KMU grant „quantumFRET“ to T.C.), the Israel Science Foundation (grants 556/22 and 3565/20 to E.L), NIH (grant R01 GM130942 to E.L. as subaward) and the Center for Nanosicence (CeNS). This work was also supported by the Federal Ministry of Education and Research (BMBF) and the Free State of Bavaria under the Excellence Strategy of the Federal Government and the Länder through the ONE MUNICH Project Munich Multiscale Biofabrication (to D.C.L.). Z.H. acknowledges a PhD scholarship from the Chinese Scholarship Council (CSC), P.D.H. the 2022-2023 Zuckerman STEM postdoctoral fellowship and P.C. and N.Z. postdoctoral fellowships from the Alexander von Humboldt foundation. T.W. acknowledges support by the Max Planck Society. We thank E. Cabrita for advice on the interpretation of NMR data.

## AUTHOR CONTRIBUTIONS

N.Z. and T.C. conceived and designed the study and supervised the project. Z.H., S.P., M.R., A.H. and D.G. performed molecular biology and produced GlnBP. Z.H, S.P., A.H. and A.N. performed biochemical characterization. Z.H. established labeling protocols. S.B. and K.J. provided SPR data. Z.H., S.P. and A.H. performed µsALEX experiments, E.B. performed PIE experiments. E.B. and D.C.L. analyzed PIE data. Z.H. and M.I. performed TIRF experiments, O.B. supported TIRF data analysis. O.B., P.D.H. and E.L. performed smFRET data analysis using mpH^2^MM. E.L. performed Gln docking calculations onto GlnBP structures. T.W. performed MD simulations. M.I., T.W. and T.C. analyzed SPR data. P.C., T.W. and T.C. analyzed and interpreted kinetic rate schemes. Z.H., P.C. and T.W. prepared figures. Z.H., P.C., T.W. and T.C. wrote the manuscript in consultation with all authors. All authors contributed to the discussion and interpretation of the results and approved the final version of the manuscript.

## MATERIAL AND METHODS

All commercially obtained reagents were used as received, unless stated otherwise. The following grades were used: Guanidine hydrochloride (99%, Sigma Aldrich), 1,4-Dithiothreit (DTT) (99%, ROTH), Thermo Scientific SnakeSkin TM Dialysis Tubing (Fisher scientific,10K MWCO, 16 mm), Ni^2+^-Sepharose resin (GE Healthcare), Albumbin fraction V (BSA), biotin-frei, ≥ 98% (Carl Roth GmbH), Imidazole, ≥ 99% (Carl Roth GmbH), Isopropyl-β -D-1-thiogalactopyranose (IPTG), ≥ 99% (Carl Roth GmbH), Kanamycin (Carl Roth GmbH), L-glutamine (Merck KGaA), L-Arginine (Carl Roth GmbH). AF555 (Jena Bioscience, Germany), AF647 (Jena Bioscience, Germany), ATTO 532 (ATTO-TEC, Germany), ATTO 643 (ATTO-TEC, Germany), mPEG3400-silane (abcr, AB111226) and biotin-PEG3400-silane (Laysan Bio Inc), Biotin-NTA (Biotium), Streptavidin (Roth, Germany), Pyranose oxidase (Sigma Aldrich, Germany), Catalase (Sigma Aldrich, Germany), Glucose (≥ 99.5% GC, Sigma Aldrich, Germany), Trolox (98%, Sigma Aldrich, Germany), Potassium hydroxide (≥85%, Honeywell, Germany), Acetone (Roth, Germany), Toluene (Roth, Germany).

### Protein expression and purification

Two GlnBP double cysteine variants were generated by site-directed mutagenesis, allowing the insertion of two cysteine residues into GlnBP at positions (V111C – G192C) and (T59C – T130C), separately. *Escherichia coli* BL21-pLysS cells were freshly transformed with the plasmid carrying the coding sequence for GlnBP WT or a GlnBP variant, and grown in 2 L LB medium (100 mg/mL Kanamycin and 50 mg/mL chloramphenicol) at 37 °C under aerobic conditions. At an OD_600nm_ of 0.6-0.8, overexpression of the proteins of interest was induced upon addition of 1 mM IPTG to the culture media. The cells were further grown for 1.5-2.0 hours after induction and then harvested by centrifugation for 20 minutes at 1,529 g (Beckman, JA10) at 4 °C. All subsequent operations were carried out at 4 °C, and all solutions were stored at 4 °C. Cell pellets from 2 L culture were collected in a 50 mL falcon and resuspended in buffer A (50 mM Tris-HCl, pH 8.0, 1 M KCl, 10 mM imidazole, 10% glycerol) with 1 mM dithiothreitol (DTT). and gently shaken overnight at 4 °C.

Cells were disrupted by sonication (Branson tip sonication; amplitude: 25%; 10 min; 0.5 s on-off pulses; temperature was kept low by the use of an ice-water bath). Centrifugation was used to fractionate the cell lysate (at 4 °C for 30 min at 4,416 g, Eppendorf, Centrifuge 5804 R) and at 4 °C for 1 hour for ultracentrifugation (70,658 g, Beckman, Type 70Ti) in vacuum, and the pellet was discarded. The protein was purified by affinity chromatography using the Ni^2+^-Sepharose fast flow resin (GE Healthcare), pre-equilibrated with 10 column volumes of buffer A containing 1 mM DTT and gravity loaded with the supernatant from the preceding ultra-centrifugation step. The resin-bound protein was washed with 10 column volumes of buffer A containing 1 mM DTT, followed by buffer B containing 1 mM DTT (50 mM Tris-HCl, pH 8,0, KCl 50 mM, imidazole 20 mM, glycerol 10%), and finally eluted in buffer C (50 mM Tris-HCl, pH 8.0, KCl 50 mM, imidazole 250 mM, glycerol 10%) with 1 mM DTT. The eluted sample was concentrated (Vivaspin6 columns, 10 kDa MWCO, 6 mg/mL), dialyzed against PBS buffer supplemented with 1 mM DTT, and stirred gently at 4 °C overnight. SDS-PAGE was used to quantify the yield of protein overexpression and purification (Comassie staining). The absorbance at 280 nm was used to estimate the protein concentration (knowing the molar extinction coefficient of GlnBP ∼25,900 M^-1^ cm^-1^). The protein was then split into aliquots and kept at −20 °C. All proteins were further purified using size-exclusion chromatography (ÄKTA pure system, Superdex 75 Increase 10/300 GL, GE Healthcare). The purified protein was split into aliquots and stored at −80 °C prior to the measurements.

### Unfolding and refolding process of GlnBP WT and GlnBP variants

The stock concentrations of GlnBP variants were estimated at about 6 mg/mL. Each GlnBP variant was thawed from −80 °C, then the protein was diluted to a final concentration of 3-4 µM (final volume of ̴20 mL) in the unfolding buffer (PBS buffer) containing 6 M guanidine hydrochloride (GndHCl). Subsequently, the solution was incubated for 3 hours under gentle stirring at ambient temperature. Next, the unfolded GlnBP variants were centrifuged (3,046 g, 30 min at 4 ℃) to remove insoluble aggregates which could act as nuclei to trigger aggregation during refolding process. A Snakeskin TM dialysis membrane was prepared (pre-cooled at 4 °C and soaked in refolding buffer - PBS buffer with 1 mM DTT, pH 7.4 - for 2 min). The GlnBP variants were transferred into the dialysis tubing, which were sealed tightly afterwards by double-knots and clips at each end. The unfolded GlnBP variant was refolded by a two-step dialysis, in the presence of a total 200-fold excess of refolding buffer. First, each protein was dialyzed against 2 L refolding buffer overnight under gentle stirring at 4 °C. Then, buffer was exchanged with additional 2 L refolding buffer for another day at 4 °C. The refolded protein was then concentrated from 20 mL to final 500 μL (Vivaspin 10 kDa MWCO; 3,000 g × 15 min at 4 °C) and further purified by size-exclusion chromatography (ÄKTA pure system, Superdex-75 Increase 10/300 GL, GE Healthcare). The unfolding and refolding process for GlnBP WT was conducted under the same conditions as described for the GlnBP variants.

### Isothermal titration calorimetry (ITC) measurements

The ITC measurements were performed in a MicroCal PEAQ-ITC isothermal titration calorimeter (Malvern Instruments). The prediction ITC software “MicroCal PEAQ-ITC Control” was employed for designing and conducting the experiments. Once the K_d_ value and the binding stoichiometry (N) were assigned as predefined values, the concentration of both the protein and the titrant (ligand) stock solutions could be calculated by the “design-experiment” function on the software to get an optimal sigmoidal one-site binding curve. GlnBP concentration was assessed using the Nanophotometer (N60 Touch, Implen GmbH) with at least three reading repeats to get accurate determinations of concentration values. For all ITC measurements, the temperature was set at 25 °C with stirring speed at 750 rev / min. The GlnBPs solution (10 μM in PBS buffer pH 7.4, 300 μL) was manually loaded into the sample cell. The titrant (L-Glutamine, 100 μM in PBS buffer, pH 7.4) was automatically loaded into the titration syringe and injected in the sample cell with a titration speed of 2 μL every 150 second and a total of 19 injections. As a control experiment, L-Glutamine was titrated into the sample cell containing PBS buffer without GlnBPs. All the titration data were analyzed using the MicroCal PEAQ-ITC Analysis Software.

### Surface plasmon resonance spectroscopy (SPR) and data analysis

SPR assays were performed on a Biacore T200 (Cytiva) using a CM5 Series S carboxymethyl dextran sensor chip coated with His-antibodies from the Biacore His-capture kit (Cytiva). Briefly, the chips were equilibrated with running buffer until the dextran matrix was swollen. Afterwards, two flow cells of the sensor chip were activated with a 1:1 mixture of N-ethyl-N-(3-dimethylaminopropyl) carbodiimide hydrochloride and N-hydroxysuccinimide according to the standard amine coupling protocol. A final concentration of 50 µg/mL anti-histidine antibody in 10 mM acetate buffer pH 4.5 was loaded onto both flow cells using a contact time of 420 s for gaining a density of approximately 10,000 resonance units (RU) on the surface. By injection of 1 M ethanolamine/HCl pH 8.0, free binding sites of the flow cells were saturated. Preparation of chip surfaces was carried out at a flow rate of 10 µL/min. All experiments were carried out at a constant temperature of 25 °C using PBS buffer (0.01 M phosphate buffer, 2.7 mM KCl, 0.137 M NaCl, pH 7.4) supplemented with 0.05 % (v/v) detergent P20 as running buffer.

For interaction analysis, GlnBP-6His (1.5 µM) was captured onto one flow cell using a contact time of 240 s at a constant flow rate of 10 µL/min. This resulted in a capture density of approximately 1,200 RU of GlnBP-6His. Eight different concentrations of glutamine (7.8, 15.6, 31.25, 62.5, 125, 250, 500 and 1,000 nM) were injected onto both flow cells using an association time of 50 s and a dissociation time of 360 s. The flow rate was kept constant at 30 µL/min. As control, running buffer was injected. The chip was regenerated after each cycle by removing GlnBP-6His completely from the surface using 10 mM glycine pH 1.5 for 60 s at a flow rate of 30 µL/min.

Sensorgrams were recorded using the Biacore T200 Control software 2.0.2. The surface of flow cell 1 was not coated with GlnBP-6His and used to obtain blank sensorgrams for subtraction of the bulk refractive index background with the Biacore T200 Evaluation software 3.1. The referenced sensorgrams were normalized to a baseline of 0. Peaks in the sensorgrams at the beginning and the end of the injection are due to the run-time difference between the flow cells for each chip.

In total, 26 SPR sensorgrams in three sets of measurements were recorded. To correct for remaining drift in the sensorgrams, the initial 60 s of the sensorgrams prior to Gln injection and the last 100 s of the dissociation phase where first fitted with an exponential function, which was subtracted from the sensorgrams. The drift-corrected sensorgrams were fitted to the reaction scheme of Eq. (1) based on the differential equations^67,106^.

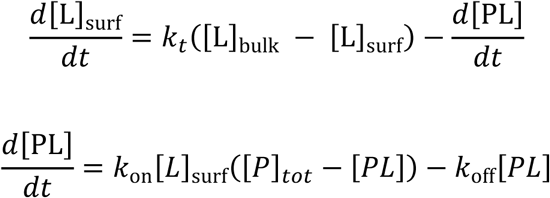

where [*L*]_bulk_ = [Gln] and [*L*]_surf_ are the free glutamine concentrations in the bulk flow and at the sensor surface, [𝑃]_tot_ is the total concentration of surface-immobilized protein, and [𝑃*L*] is the concentration of bound protein complexes. Conversion to the SPR binding response *r* via [𝑃*L*] = α *r* and [𝑃]_tot_ = α *r*_𝑚𝑎𝑥_ leads to fit results for the binding rate constants that are insensitive to the (unknown) conversion factor α, which can be understood from the fact that the quasi-steady-approximation *d*[*L*]_surf_⁄*d*𝑡 ≈ 0 holds for SPR setups^67,106^. The association phases of the sensorgrams were fitted with initial conditions [*L*]_𝑠𝑢*r*𝑓_ = 0 and *r* = 0 and fit parameters *k_t_* and *r*_𝑚𝑎𝑥_at different values of *k*_on_ after substitution of *k*_off_ by *K*_d_ *k*_on_. Prior to these fits with fixed *k*_on_, a remaining small vertical off-set of the sensorgrams was determined as additional fit parameter in fits with unconstrained, large *k*_on_ and subtracted from the sensorgrams. The first 50 s of the dissociation phases were fitted with single fit parameter *k_t_* for the initial conditions [*L*]_𝑠𝑢*r*𝑓_= [*L*]_𝑏𝑢𝑙*k*_and *r* = *r*_𝑚𝑎𝑥_⁄(1 + *K*_*d*_⁄[*L*]_𝑏𝑢𝑙*k*_), with *r*_𝑚𝑎𝑥_determined from fits of the association phase of the sensorgram for unconstrained, large *k*_on_. Background-corrected sensorgrams that do not reach binding equilibrium in the association phase (because of small [Gln]), still show marked drifts in binding equilibrium, or do not resolve the initial increase of the binding signal of the association phase (because of large [Gln]) were discarded, which leads to the 13 sensorgrams of Figures 6B,C and Figure 6—figure supplement 1with fit results for α = 1 μM/RU. Fits with e.g., α = 1 mM/RU (not shown) lead to practically identical results. All fits were conducted with Mathematica 13 based on the functions ParametricNDSolveValue to obtain numerical solutions of the differential equations and NonlinearModelFit for fitting parameters of these solutions.

### Protein labeling

The refolded GlnBP(111C-192C) and GlnBP(59C-130C) variants were labeled with commercial maleimide derivatives of AF555/AF647 or ATTO 532/ATTO 643^102^, and then purified by SEC. The chromatogram of refolded GlnBP(111C-192C) labeled with AF555/AF647 is shown in Figure 2B, and those of all other variants and dye labeling combinations are displayed in Figure 2—figure supplement 2. First, the His-tagged protein was incubated in 10 mM DTT in PBS buffer for 30 min to reduce all oxidized cysteine residues. Subsequently, the protein was diluted 10 times with PBS buffer and immobilized on a Nickel Sepharose 6 Fast Flow resin (GE Healthcare). The resin was washed extensively with milliQ water followed by PBS buffer pH 7.4. To remove the excess of DTT, the resin was washed with PBS buffer. The protein was left on the resin and incubated overnight at 4 °C with 5-10 times molar dye excess in PBS buffer pH 7.4. Subsequently, the unreacted fluorophores were removed by washing the resin with 6 mL of PBS buffer. Bound proteins were eluted with 800 μL of elution buffer (PBS buffer, pH 7.4 400 mM Imidazole) The labeled protein was further purified by size-exclusion chromatography (ÄKTA pure, Superdex-75 Increase 10/300 GL, GE Healthcare) to eliminate remaining fluorophores and remove other contaminants and soluble aggregates. The selected elution fractions were used without further treatment for smFRET experiments as described below. In general all experiments were carried out at room temperature using 25–50 pM of double-labeled GlnBP protein in PBS buffer (pH7.4). Titration experiments were performed by adding specific concentrations of ligand (glutamine) to the buffer.

### smFRET experiments with µsALEX

Single-molecule μsALEX experiments were carried out at room temperature on a custom-built confocal microscope. In short, alternating excitation light (50 μs period) was provided by two diode lasers operating at 532 nm (OBIS 532-100-LS, Coherent, USA) and 640 nm (OBIS 640-100-LX, Coherent, USA). Both lasers were combined by coupling them into a polarization maintaining single-mode fiber (P3-488PM-FC-2, Thorlabs, USA) and subsequently guided into the microscope objective (UplanSApo 60X/1.20W, Olympus, Germany) via a dual-edge dichroic mirror (ZT532/640rpc, Chroma, USA). In general, the 532 and 640 nm diode lasers operated at 60 and 25 μW, respectively (measured at the back aperture of the objective), unless stated otherwise. Fluorescence light was collected by the same objective, focused onto a 50 μm pinhole and separated into two spectral channels (donor and acceptor fluorescence) by a dichroic beamsplitter (H643 LPXR, AHF, Germany). Fluorescence emission was collected by two avalanche photodiodes (SPCM-AQRH-64, Excelitas) after additional filtering (donor channel: BrightLine HC 582/75 and acceptor channel: Longpass 647 LP Edge Basic, both from Semrock, USA). The detector outputs were recorded via an NI-Card (PCI-6602, National Instruments, USA) using a custom-written LabView program.

### smFRET data analysis (µsALEX)

Data analysis for µsALEX was performed using an in-house written software package as previously described^16^. Three relevant photon streams were extracted from the recorded data based on the alternation period, corresponding to donor-based donor emission F(DD), donor-based acceptor emission F(DA) and acceptor-based acceptor emission F(AA). Bursts from single-molecules were identified using published procedures^59^ based on an all-photon-burst-search algorithm with a threshold of 15, a time window of 500 μs, and a minimum total photon number (F(DD)+D(DA)+F(AA)) of 150, unless stated otherwise in the figure caption.

For each fluorescence burst, the stoichiometries S* and apparent FRET efficiencies E* were calculated and then presented for all bursts yielding a two-dimensional (2D) histogram. Uncorrected apparent FRET efficiency, E*, monitors the proximity between the two fluorophores and is calculated according to E* = F(DA)/(F(DD)+F(DA)). Apparent stoichiometry, S*, is defined as the ratio between the overall fluorescence intensity during the green excitation period over the total fluorescence intensity during both green and red periods and describes the ratio of donor-to-acceptor fluorophores in the sample: S*=(F(DD)+F(DA)/(F(DD)+F(DA)+F(AA)). Collecting the E* and S* values of all detected bursts into a 2D E*-S* histogram yielded subpopulations that can be separated according to their E*- and S*-values. The 2D histograms were fitted using a 2D gaussian function, yielding the mean apparent FRET efficiency and its standard deviation or width of the distribution. µsALEX, assists in sorting single molecules based on their donor/acceptor dye brightness ratio (stoichiometry S*) and uncorrected mean FRET efficiency (apparent FRET E*), which can be related on the mean inter-dye distance^102,107^.

Analysis with mpH^2^MM was conducted as described previously by the Lerner lab^62^. In short, the FRET Bursts software^108^ was used for detecting single-molecule photon bursts using the dual channel burst search^59^ AND-gate algorithm with a sliding window of m=10 photons searching for instances with an instantaneous photon rate of at least F=6 times the background rate. Afterwards, bursts of such consecutive photons were filtered to have at least 50 photons originating from donor excitation and at least 50 photons originating from acceptor excitation. In the data analysis, the photon stream was then divided into photon streams of different bursts, and a time shift was applied to acceptor excitation originating photons stream so that their arrival time range overlap with that of donor excitation originating photon streams. Optimizations were conducted with state models of increasing numbers of states, and the *Viterbi* algorithm was employed for calculating the integrated complete likelihood (ICL). Optimizing for larger numbers of states ceased once the ICL ceased to decrease between successively larger state models. Optimized models were manually examined, and the optimal state model selected considering the ICL and the reasonableness of the model given prior knowledge based on transition rates and the E* and S* values of the states. After selection of the most-likely state model, the corresponding most-likely state-path determined by the *Viterbi* algorithm was used to segment bursts into dwells and to classify burst by which states were present within each burst.

To support the idea that apo and holo state in solution match with that of the crystal structure, we performed a quantitative comparison of inter-dye distances calculated from dye accessible volumes (AV) on structural models of apo and holo protein, and those derived from the experimental smFRET results. For dye AV calculations, we used the FPS method, established by the Seidel lab^109^ (Figure 2—figure supplement 1). The experimental data were corrected for setup-dependent parameters according to refs.^58,103^ to obtain accurate FRET values from µsALEX data. Using a Förster distance of 5.2 nm for AF555/AF647, we found good agreement, i.e., 0.3-0.5 nm deviations (and 1.0 nm in one case) between the calculated and experimentally derived inter-dye distances for both mutants (Figure 2—figure supplement 1).

### smFRET measurements with MFD-PIE and burst-wise FCS analysis

Solution-based smFRET experiments were performed on a home-built dual-color confocal microscope that combines multiparameter fluorescence detection (MFD) with pulsed interleaved excitation (PIE)^65^. MFD-PIE experiments have been described in detail previously^110^. With MFD-PIE, it is possible extract FRET efficiency, stoichiometry, fluorescence lifetime and fluorescence anisotropy information from each single-molecule burst. Correction factors including direct acceptor excitation (α), spectral crosstalk (β) and detection correction factor (γ) are also accounted for reporting accurate the FRET efficiency values^111^. The accurate FRET efficiency (E) can be determined from:

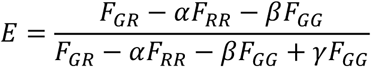

where F_GG_, F_GR_ and F_RR_ are background-corrected fluorescence signals detected in green/ donor (G), red/acceptor (R) after donor excitation and acceptor channels, respectively.

Alternatively, the use of picosecond pulsed lasers and time-correlated single photon counting (TCSPC) electronics enable calculating FRET efficiencies from the quenching of the donor in presence of acceptor. According to the formula:

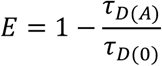

𝜏_𝐷(𝐴)_ is the fluorescence lifetime of the donor in presence of acceptor and 𝜏_𝐷(0)_ is the fluorescence lifetime of the donor only species. Static species can be observed on the theoretical static FRET line, which is a linear relation between E and 𝜏_𝐷(𝐴)_. Sub-ms conformational dynamics can also be identified and judged by observing the right-shifted populations from the static FRET line.

For the measurements here, 100 pM of GlnBP labeled with ATTO 532 and ATTO 643 was placed on a BSA-passivated LabTek chamber and measured for 2 hours. The sample was excited with 532 and 640 nm pulsed lasers with a repetition rate of 26.6 MHz and laser powers of 45 and 23 µW (measured at the back aperture of the objective), respectively.

Burst-wise FCS analysis is an alternative approach to observe sub-ms conformational dynamics. In this approach, donor (DD) and acceptor (AA) signals detected from single-molecule events are cross-correlated. Thus, fluctuations in the FRET efficiencies appear as an anti-correlated signal in the donor-acceptor fluorescence cross-correlation function. Burst with sufficient photons detected in both the donor and acceptor channels were selected. A time window of 50 ms was applied around each burst. If another burst was detected within this time window, both were eliminated to ensure correlation functions that are specific to the selected bursts. All the above mentioned data analysis was done using the PIE analysis with Matlab (PAM) software package^112^.

### Surface immobilization of DNA and GlnBP(111C-192C)

Biotin-streptavidin interaction was used to immobilize tagged proteins and labeled DNA on a PEG-functionalized coverslip for single molecule studies. The protein-his-tag and a biotin-NTA chelated with Ni^2+^ were used to mark GlnBP(111C-192C) labeled with maleimide modified derivatives of ATTO 532/ATTO 643, whilst DNA labeled with Cy3B/ATTO 647N was directly tagged with a biotin. To prepare a functionalized glass surface, cover slides (1.5H Marienfeld Superior) were first sonicated in MQ water for 30 min. The slides were rinsed three times with MQ water, sonicated for 30 min in HPLC-grade acetone, rinsed three times with MQ water again. Then, the slides were sonicated with 1 M KOH for 30 min, rinsed three times with MQ water and dried with a stream of nitrogen air. To remove any organic material left on the surface, the cover slides were plasma-cleaned for 15 min with oxygen. To create a mPEG/biotin–coated surface, the slides were immediately incubated in a 99:1 solution of mPEG3400-silane (abcr, AB111226) and biotin-PEG3400-silane (Laysan Bio Inc) in a Toluene solution overnight at 55 °C. After incubation, the slides were sonicated (10 min in ethanol, 10 min in MQ water), dried under nitrogen stream, and kept under vacuum. Prior to TIRF experiments, each slide was incubated with a 0.2 mg/mL streptavidin in PBS solution for 10 min utilizing Ibidi sticky-slide (18 well) for single molecule studies. PBS buffer pH 7.4 was used to wash away the unbound excess of streptavidin. For GlnBP(111C-192C) immobilization, 20 nM biotin-NTA (QIAGEN) was charged with 50 nM Ni^2+^ and incubated on the slide for 10 min before rinsing away the unbound excess biotin-NTA and Ni^2+^ with PBS (this step was omitted for the labeled DNA samples). GlnBP(111C-192C) at 0.8 nM and dsDNA at 0.04 nM were incubated for 5 and 1 min, respectively. For single-molecule data collection, imaging buffer (PBS, pH 7.4) containing 2 mM Trolox was used. For dsDNA we used PBS buffer in combination with an oxygen scavenging system (pyranose oxidase at 3 U/mL, catalase at final concentration of 90 U/mL, and 40 mM glucose). After that, the chambers were sealed with Silicone Isolators^TM^ Sheet Material (Grace Bio-labs). All the single-molecule investigations were done at room temperature.

### smFRET measurements with TIRF microscopy including data analysis

Single-molecule TIRF measurements were conducted on a homebuilt microscope using an Olympus iX71 inverted microscope body. Light from a 532 nm continuous wave laser (532 nm OBIS, Coherent) was transmitted off-axis onto the back-focal plane of a microscope objective (UAPON TIRF 100X 1.49NA, Olympus) via a dualband dichroic beamsplitter (TIRF Dual Line Beamsplitter zt532/640rpc, AHF Analysetechnik) to generate total internal reflection at the glass-water interface. Fluorescent emission was then split spectrally using a Dual View System (DV2, Photometrics) equipped with a dichroic beamsplitter (zt640rdc, AHF Analysetechnik). The two emission channels were then spectrally filtered using emission filters (582/75 Brightline HC and 731/137 BrightLine HC respectively, both AHF Analysetechnik). Image series were acquired using an EMCCD camera (C9100-13, Hamamatsu) in combination with the µManager^108^ software. The iSMS^109^ software was used to retrieve and calculate traces of the donor and acceptor fluorescence intensity from consecutive fluorescent images.

### MD simulations

Starting point of our atomistic simulations with the AMBER20 software package^69^ and the ff14SB force field parameters^71^ was the ligand-bound crystal structure with PDB identifier 1WDN. The protein state of the titratable amino acids and including ligand was determined with the software PROPKA3^113^ The protein, with and without ligand, was solvated in explicit TIP3P water in a octahedral simulation box with a minimum distance of 15 Å of protein atoms to the box boundaries at a salt concentration of 150 mM. The two simulation systems with and without ligand were carefully relaxed in 9 steps according to the AMBER tutorial “Relaxation of explicit water systems” (see https://ambermd.org/tutorials/). Production simulations starting from the system conformations obtained after relaxation were performed with the AMBER20 software implementation for graphics processing units (GPUs)^69,70^ with a time step of 4 fs after hydrogen-mass repartitioning^114^. In these simulation, the temperature was kept at 300 K using a Langevin thermostat with a collision frequency of 1 ps^-1^ and the pressure was kept at 1 bar with the Berendsen barostat.

To further investigate the coupling between conformational changes of GlnBP and ligand binding/unbinding, we performed atomistic simulations starting from the ligand-bound GlnBP structure with the AMBER20 software implementation for graphics processing units (GPUs)^69,70^ and the ff14SB force field parameters^71^ (see Methods for details). To observe unbinding events on the microsecond timescale accessible in our simulations, we reduced all interactions between the protein and the ligand by 16% by rescaling the partial charges and ε parameters of the ligand atoms with the commands *change charge* and *changeLJSingleType* <u>of the program ParmEd implemented in Amber.</u>

## Supporting information for Han et al

### Supplementary Figures

**Figure 2—figure supplement 1.**
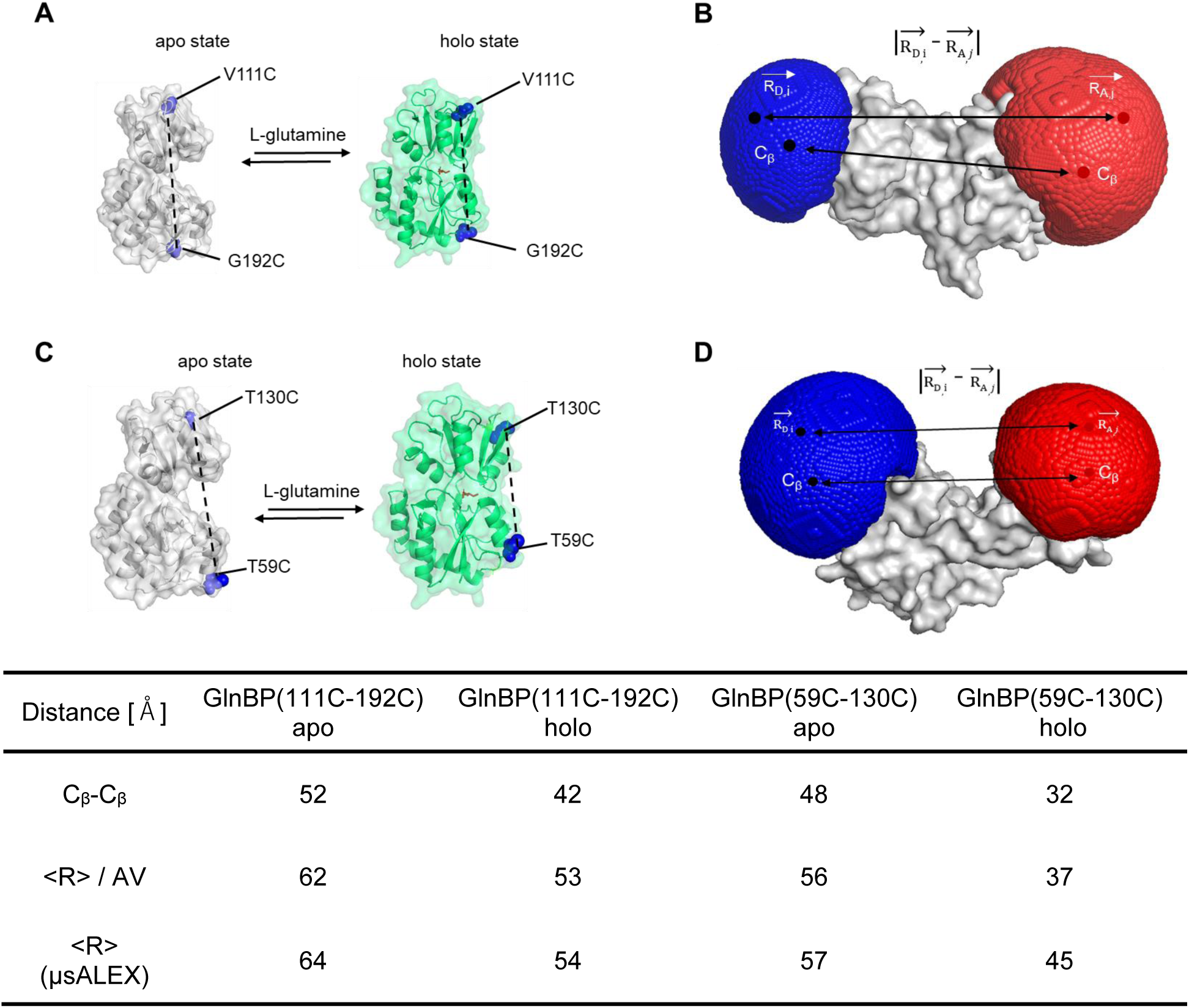
Crystal structure and dye accessible volume calculations of GlnBP cysteine variants. (A, C) Crystal structure of the ligand-free (grey structure) and ligand-bound (green structure) GlnBP with the two labeling positions of the respective variants indicated in blue. (B, D) Simulation of accessible volumes for AF555 and AF647 with values of interprobe distinces based on structural predictions (C_ß_-C_ß_ distances and fluorophore accessible volumes) and experimental values <R>.

**Figure 2—figure supplement 2.**
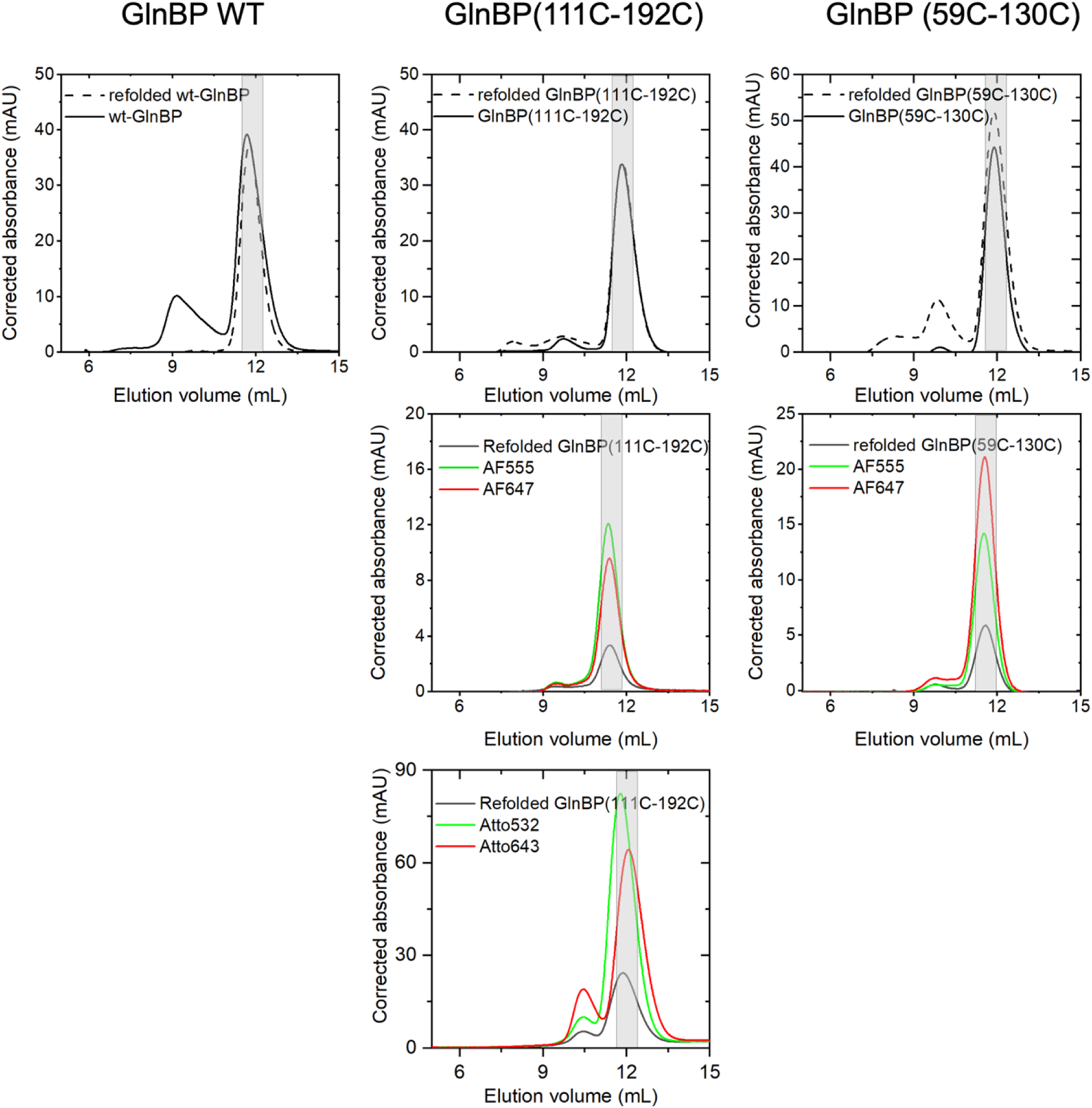
Size Exclusion Chromatography (SEC) of refolded GlnBP WT and GlnBP variants. GlnBP WT and GlnBP double-cysteine variants were unfolded with 6 M Guanidine Hydrochloride and then refolded via dialysis over two days in PBS buffer (pH 7.4, 1 mM DTT). The selected fractions (grey-shaded area) were collected and used for ITC experiments. For the solution-based smFRET measurements, the selected fractions (grey-shaded area) having the best overlap of protein, donor, and acceptor absorption were used. The protein absorption was measured at 280 nm (black curves) and the donor dye (AF555) absorption at 555 nm or donor dye (ATTO 532) absorption at 532 nm. The acceptor dye absorption (red lines) was measured at 647 nm for AF647 and 643 nm for ATTO 643.

**Figure 2—figure supplement 3.**
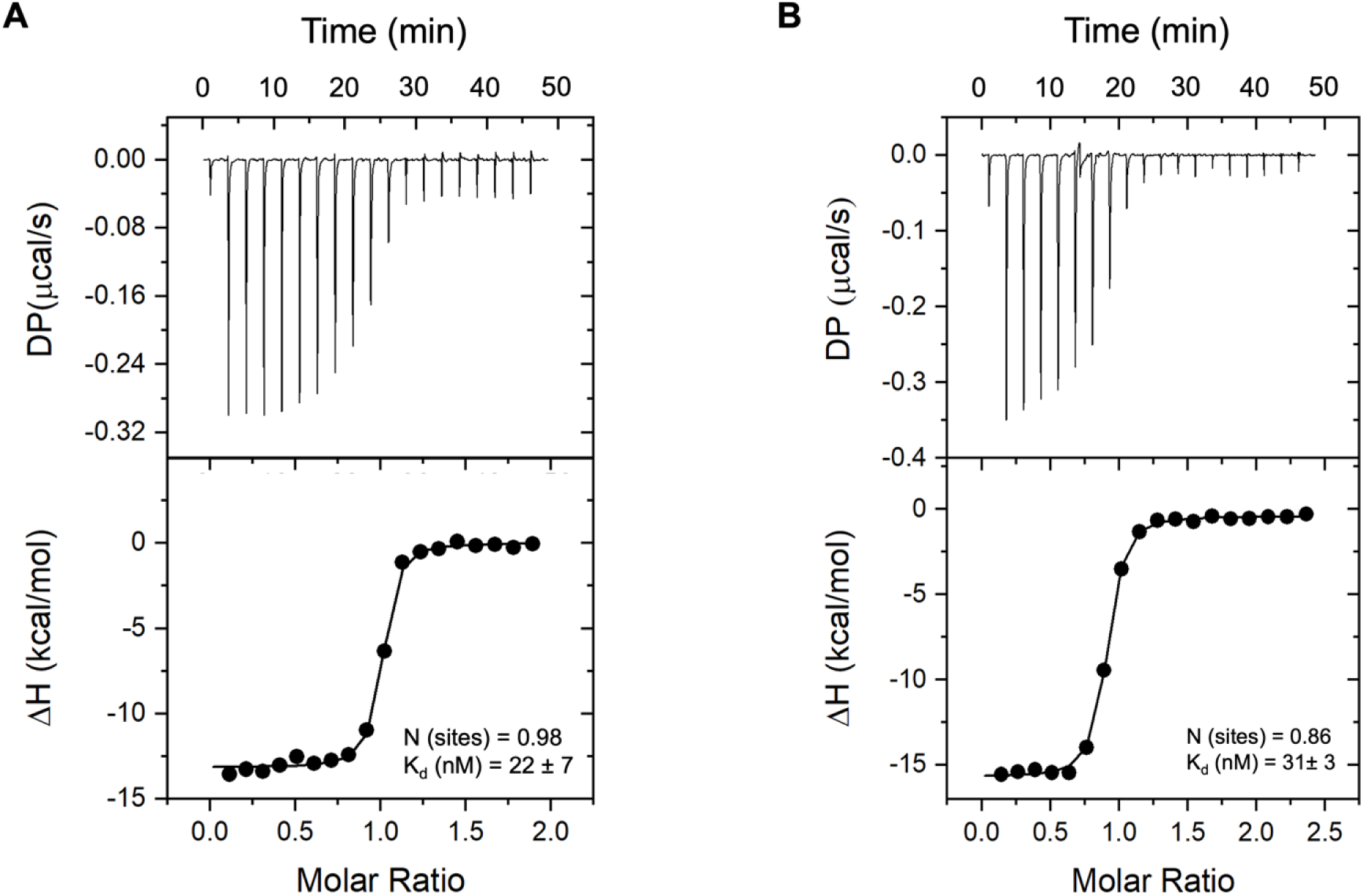
Investigating binding affinity of refolded GlnBP WT and refolded GlnBP(59C-130C) using Isothermal Titration Calorimetry (ITC) measurements. The graphs depict the changes in heat (DP, top) and enthalpy (ΔH, bottom), due to each injection of L-glutamine into the sample cell, as function of time (top x-axis of each graph) and molar ratio of refolded protein and ligand (bottom x-axis), separately. All ITC experiments were repeated three times and performed without fluorophore labeling. (A) The mean binding affinity of the refolded GlnBP WT is 22 ± 7 nM and the binding stoichiometry is close to 1. (B) The mean binding affinity of the refolded GlnBP(59C-130C) is 31 ± 3 nM and the binding stoichiometry is close to 1.

**Figure 3—figure supplement 1.**
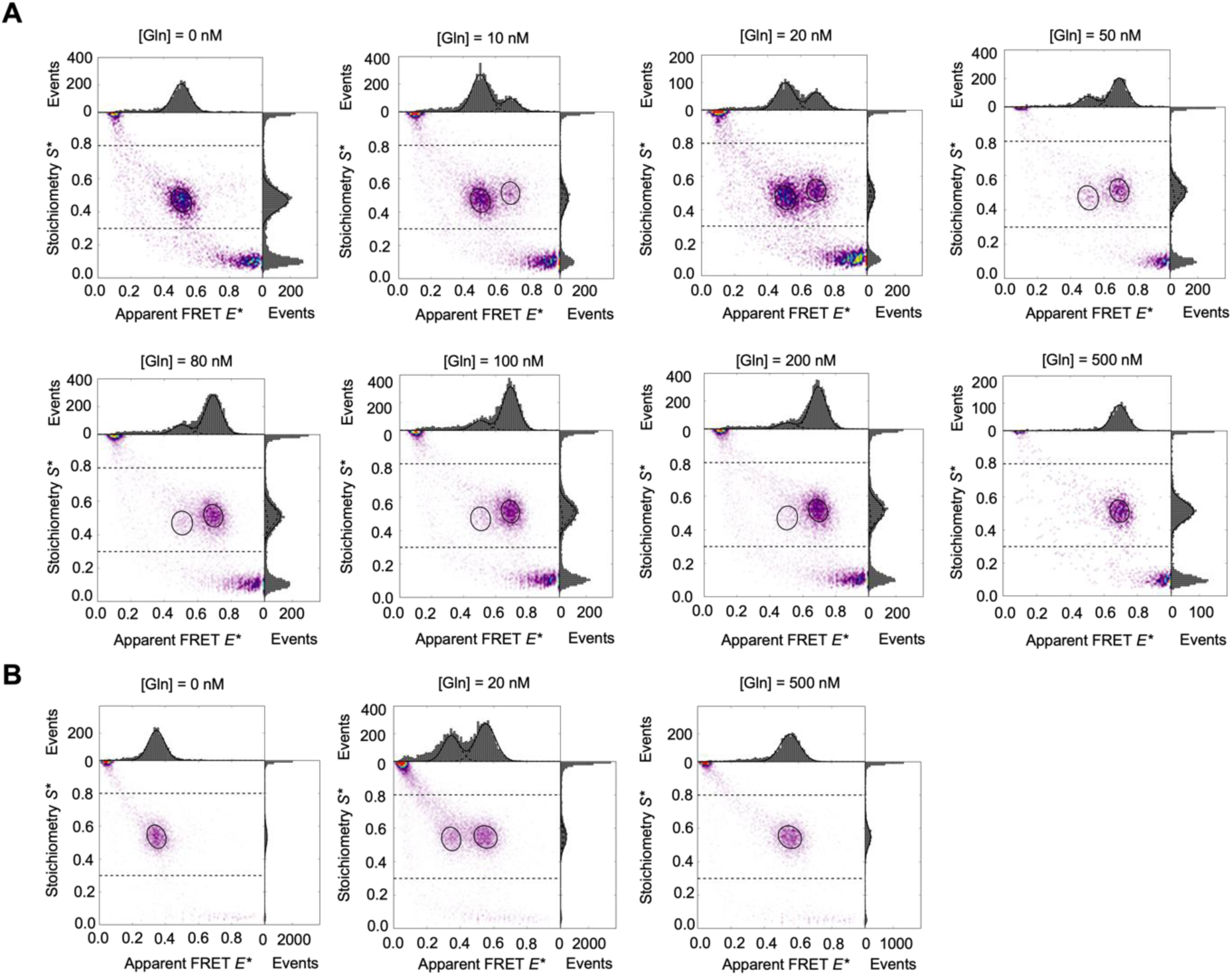
L-glutamine-induced conformational changes in refolded GlnBP(111C-192C) visualized by μsALEX measurements. μsALEX-based *E\**-*S\** histograms of the refolded GlnBP(111C-192C) double-cysteine mutants labeled with AF555/AF647 fluorophore pair (A) and labeled with ATTO 532/ATTO 643 fluorophore pair (B). First, the histograms of the apo (no L-glutamine) and holo (500 nM L-glutamine) states of the protein were fitted using a 2D gaussian distribution. Subsequently, these two distributions with variable amplitude were used to fit the intermediate ligand concentrations. Refolded GlnBP(111C-192C) labeled with AF555/AF647 shows an open state at *E\** = 0.507 and a closed high-FRET state at *E\** = 0.694 in the presence of a saturating concentration of L-glutamine. Refolded GlnBP(111C-192C) labeled with ATTO 532/ATTO 643 shows an open state at *E\** = 0.346 and a closed high-FRET state at *E\** = 0.552 in the presence of a saturating concentration of L-glutamine.

**Figure 3—figure supplement 2.**
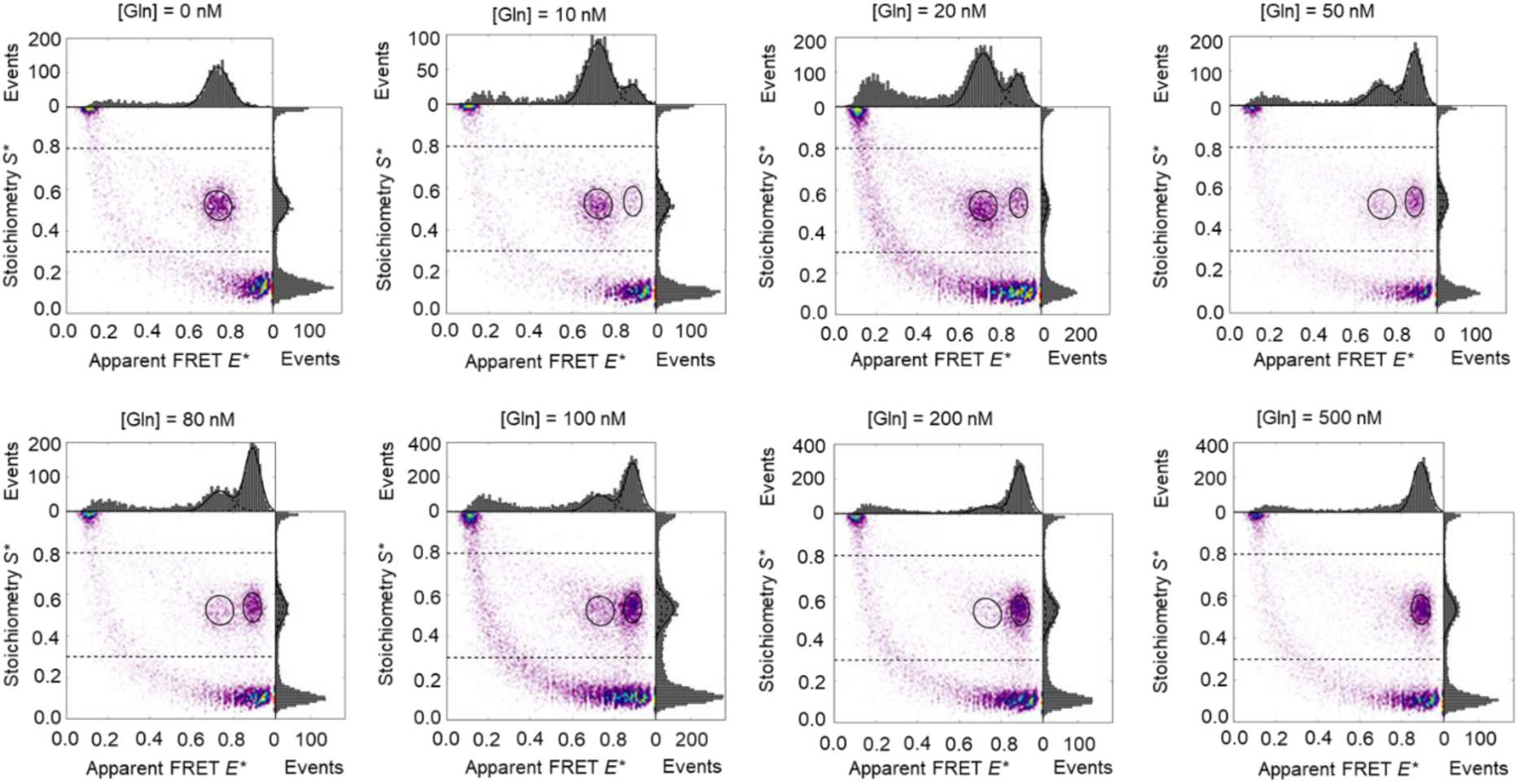
L-glutamine-induced conformational changes in refolded GlnBP(59C-130C) visualized by μsALEX measurements. μsALEX-based *E\**-*S\** histograms of the refolded GlnBP(59C-130C) double-cysteine mutants labeled with AF555/AF647 fluorophore pair. First, the histograms of the apo (no L-glutamine) and holo (500 nM L-glutamine) states of the protein were fitted using a 2D gaussian distribution. Subsequently, these two distributions were used to fit the intermediate ligand concentrations. Refolded GlnBP(59C-130C) shows an open state at *E\** = 0.735 and a closed high-FRET state at *E\** = 0.891 in the presence of a saturating concentration of L-glutamine.

**Figure 3—figure supplement 3.**
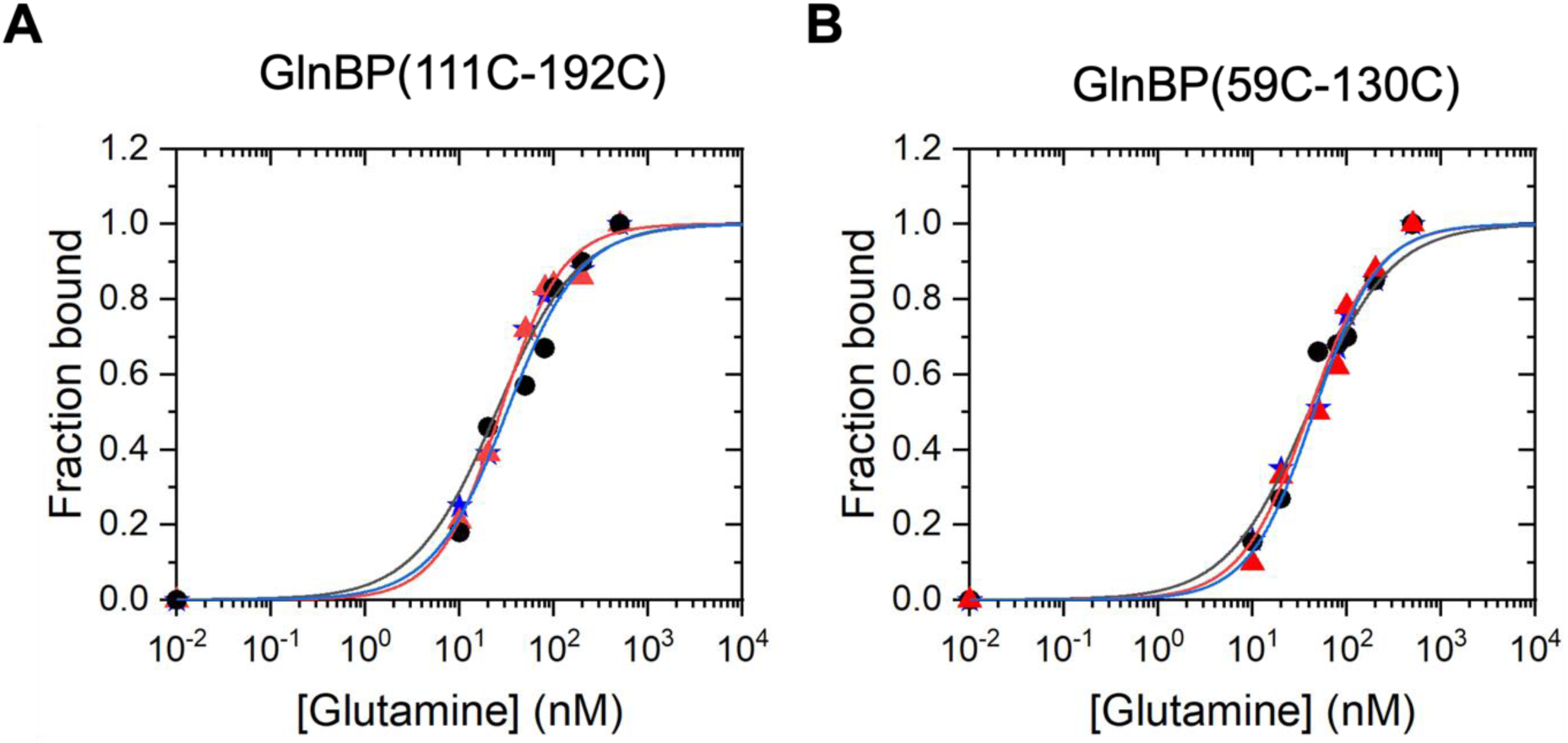
Investigating biding affinities of fluorescently labeled GlnBP variants using smFRET measurements. Binding curves of GlnBP in semilogarithmic fashion of [Gln] vs. bound fraction of protein from µsALEX experiments using the AF555/AF647 dye pair (see Fig. S3A and Fig. S4). The fraction closed, i.e., the fraction of liganded protein, was determined from the ratio of the area of the high-efficiency peak and the total peak area from the projections in the apparent FRET efficiency. The fraction bound as a function of ligand (L-glutamine) concentration was fitted with the Hill equation using Origin 2016 (Origin Lab Corp, Northampton, MA), with the maximum number of binding sites fixed to 1. All the measurements were repeated three times.

**Figure 3—figure supplement 4.**
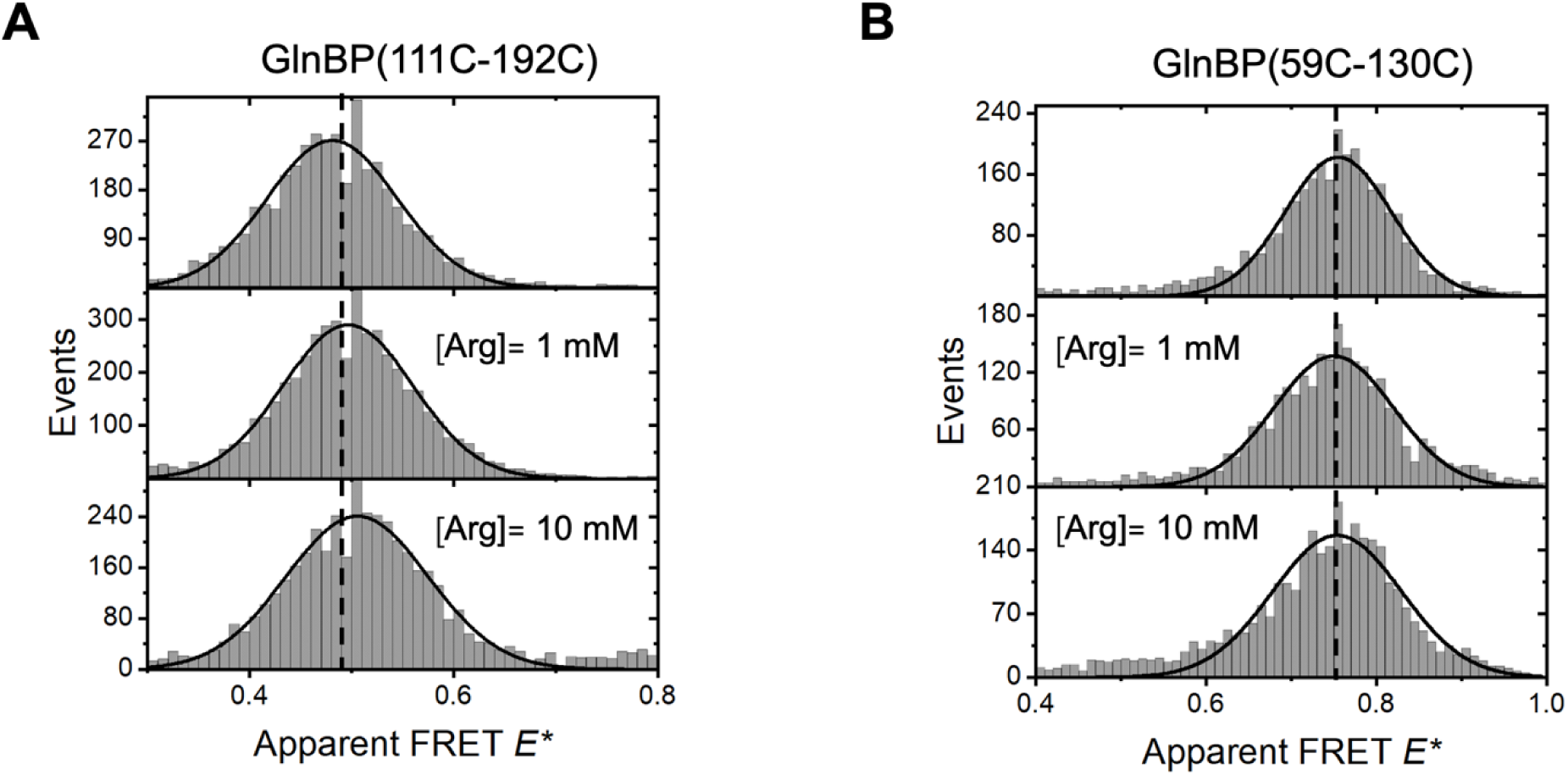
Conformational states of refolded GlnBP variants probed by solution-based μsALEX measurements reveal nearly unchanged conformations. (A) Apparent FRET efficiency histograms of refolded GlnBP(111C-192C) labelled with AF555/647 in the absence (first row) and presence of L-arginine. (B) Apparent FRET efficiency histograms of refolded GlnBP(59C-130C) labelled with AF555/AF647 in the absence (first row) and presence of L-arginine.

**Figure 3—figure supplement 5.**
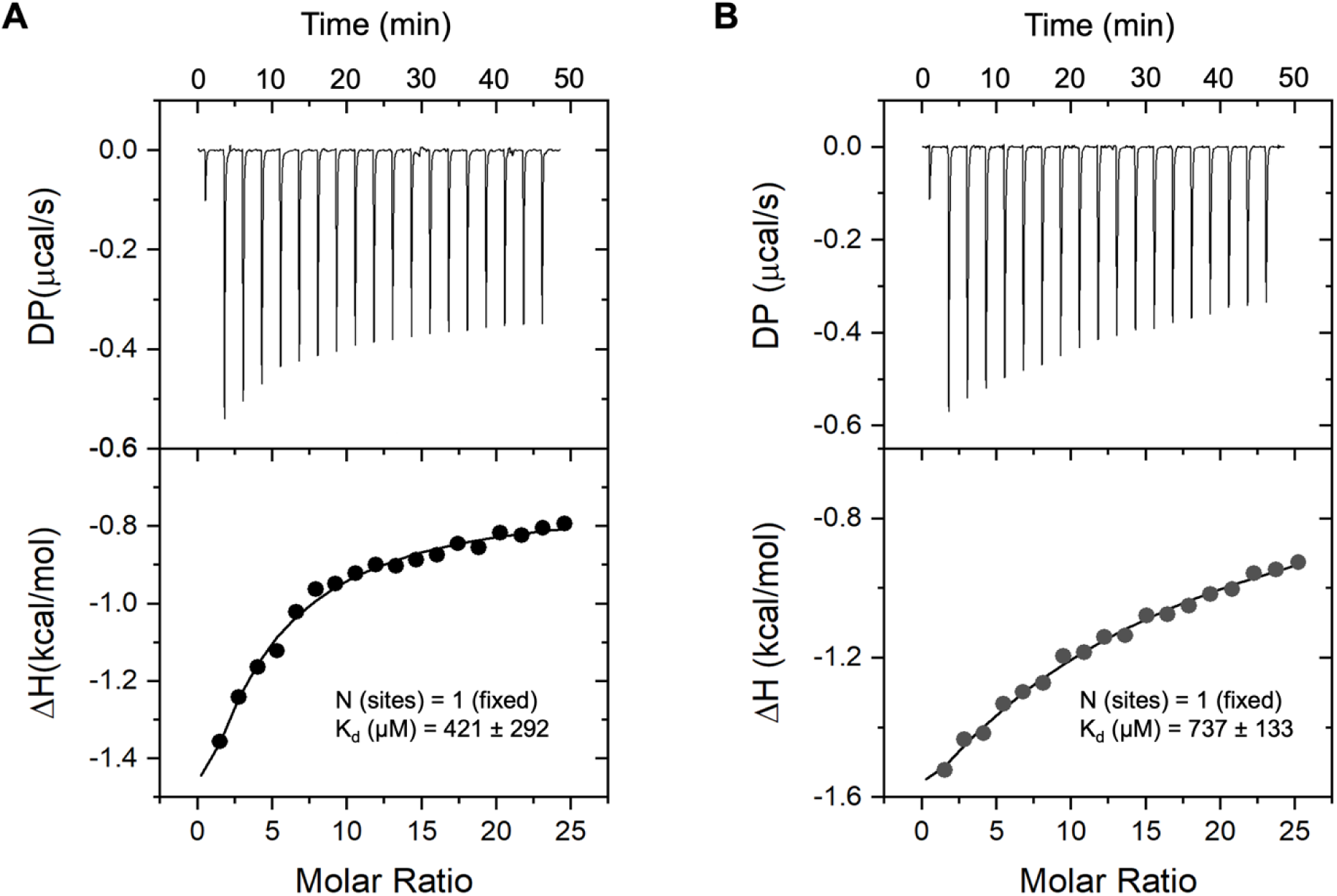
Investigating L-Arginine binding affinity of refolded GlnBP(111C-192C) and GlnBP(59C-130C) variants using Isothermal Titration Calorimetry (ITC) measurements. The graphs depict the changes in heat and enthalpy with the injection of the L-Arginine against the time and molar ratio of refolded protein and ligand, separately. All ITC experiments were repeated three times and performed without fluorophore labeling. (A) The average binding affinity of the refolded GlnBP(111C-192C) is 421 ± 292 μM. (B) The average binding affinity of the refolded GlnBP(59C-130C) is 737 ± 133 μM. The binding ratio (sites) was manually fixed to N = 1.

**Figure 4—figure supplement 1.**
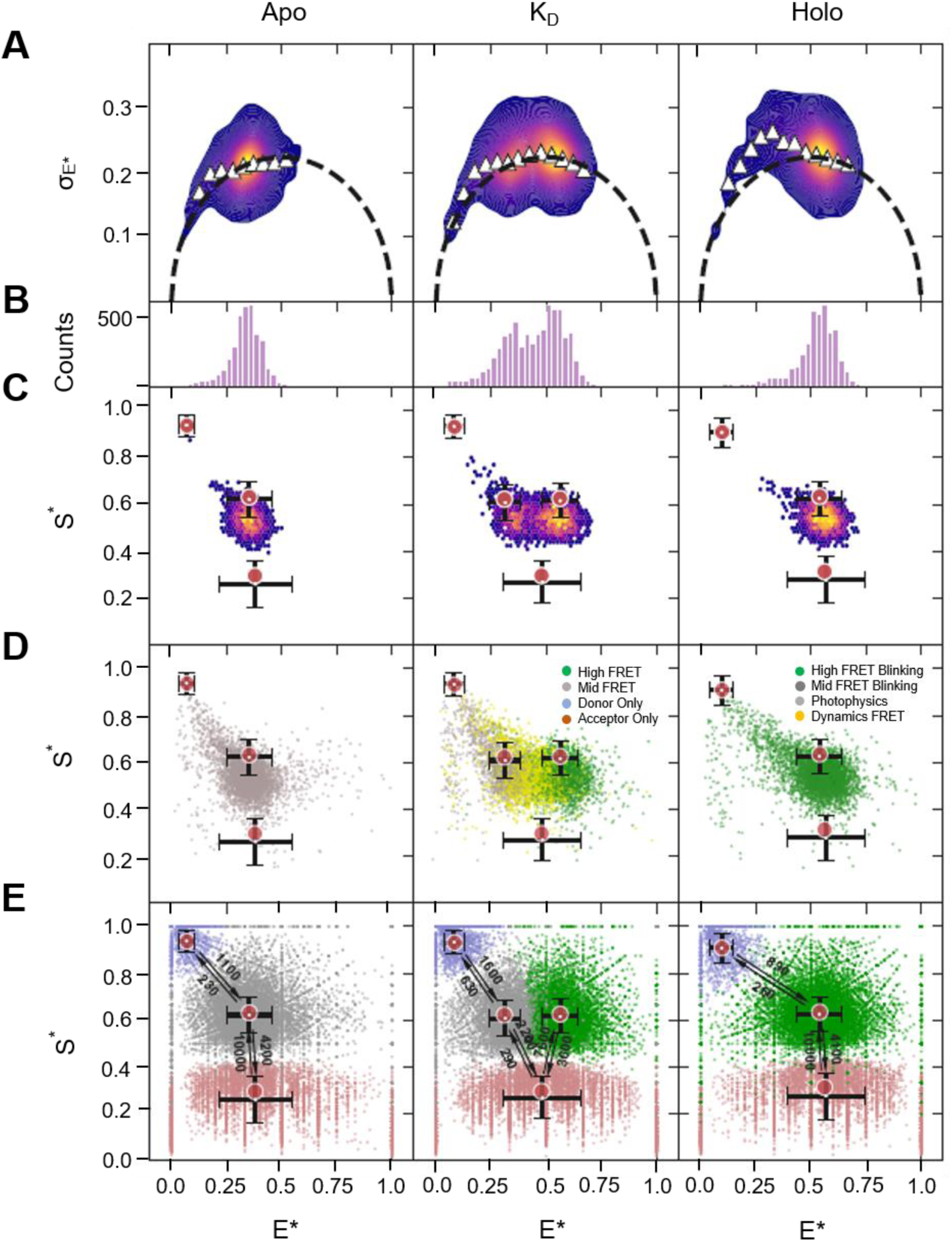
**Screening GlnBP(111C-192C) for rapid within-burst FRET dynamics**. Confocal-based single-molecule FRET results for GlnBP(111C-192C) doubly-labeled with ATTO 532 and ATTO 643, in the apo state (left panels), near the K_d_ (middle), and in the holo state (right). (A) Burst Variance Analysis (BVA) showing a weak signature of within-burst FRET dynamics in the low E* regime. (B) Histograms of E* values of bursts, (C) E* versus S* 2D histograms of bursts, (D) 2D scatter plots of bursts classified by mpH^2^MM, with colors corresponding to which state(s) are present within the bursts as determined with the Viterbi algorithm. Locations of states are given by red circles, and black crosses represent the SD of E* and S* values of dwells within each state. (E) E* versus S* 2D scatter plots of dwells in mpH^2^MM-detected states within bursts detected by the Viterbi algorithm. Red circles and black crosses are same as in (D). Arrows and adjacent numbers indicate transition rates in s^-1^ units. Transitions with rates less than 100 s^-1^ are omitted, since such slow transitions are improbable to occur within single-molecule bursts with durations shorter than 10 ms and are most probably a mathematical outcome of the mpH^2^MM framework. The dispersion of the E* and S* values of dwells in mpH^2^MM-detected states are due to the short dwell times in these states, where the shorter the dwell time in a state is, the lower the number of photons it will include, and hence the larger the uncertainty will be in the calculation of E* and S* values of dwells. E* and S* are E* and S* values uncorrected for background, since in mpH^2^MM all photons within bursts are taken into account, including ones that might be due to background.

**Figure 4—figure supplement 2.**
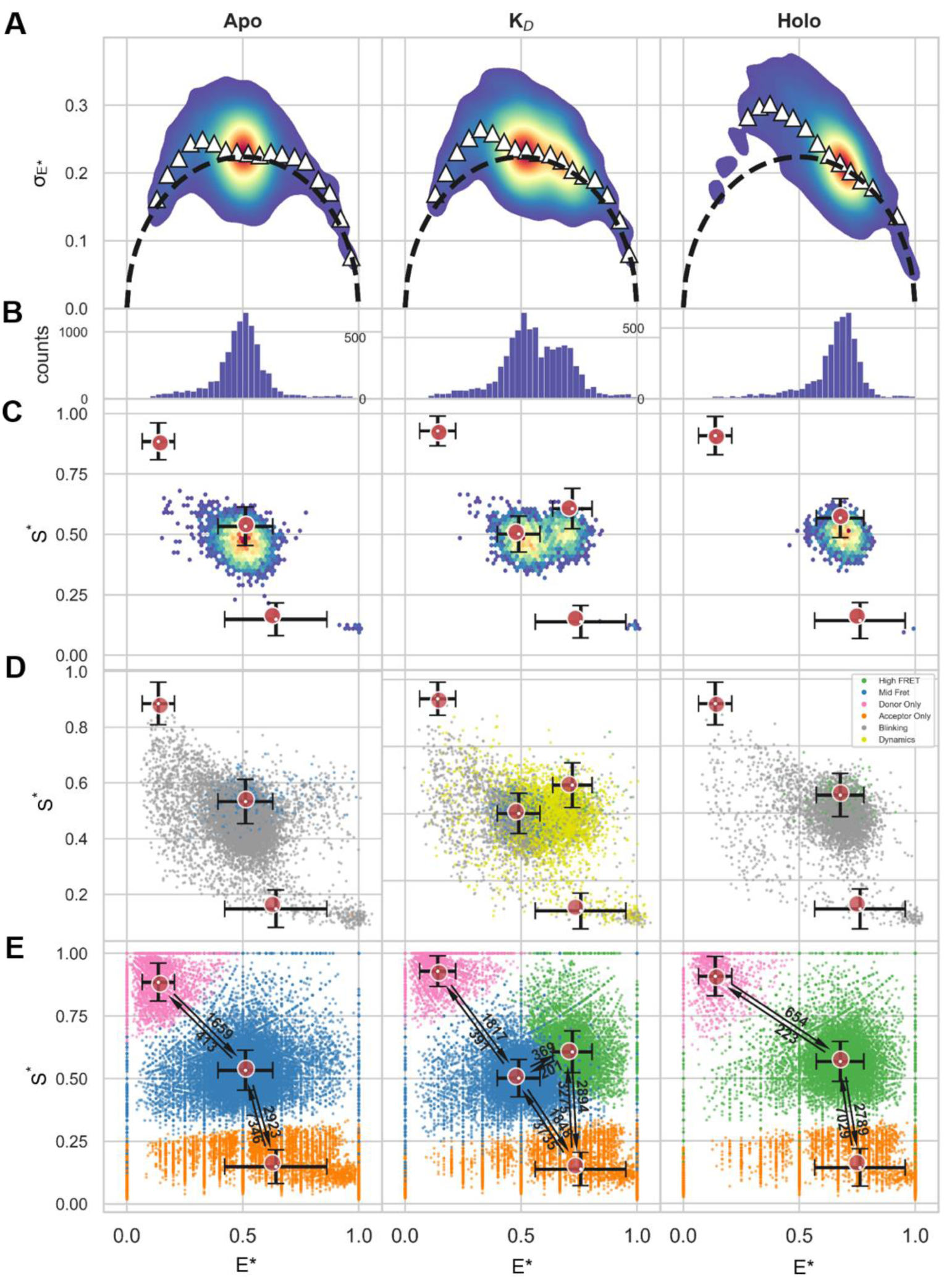
**Screening GlnBP(111C-192C) for rapid within-burst FRET dynamics**. Confocal-based single-molecule FRET results for GlnBP doubly-labeled at residues 111 and 192 with AF555 and AF647, in the apo state, near the K_D_, and holo state. (A) Burst variance analysis showing a weak signature of within-burst FRET dynamics. (B) Histograms of E* values of bursts, (C) E* versus S* 2D histograms of bursts, (D) 2D scatter plots of bursts classified by mpH^2^MM, with colors corresponding to which state(s) are present within the burst as determined with the Viterbi algorithm. Locations of states are given by red circles, and black crosses represent the SD of E* and S* values of dwells within each state. (E) E* versus S* 2D scatter plots of dwells in mpH^2^MM-detected states within bursts detected by the Viterbi algorithm. Red circles and black crosses are same as in (D). Arrows and adjacent numbers indicate transition rates in s^-1^ units. Transitions with rates less than 100 s^-1^ are omitted, since such slow transitions are improbable to occur within single-molecule bursts with durations shorter than 10 ms and are most probably a mathematical outcome of the mpH^2^MM framework. The dispersion of the E* and S* values of dwells in mpH^2^MM-detected states are due to the short dwell times in these states, where the shorter the dwell time in a state is, the lower the number of photons it will include, and hence the larger the uncertainty will be in the calculation of E* and S* values of dwells. E* and S* are E* and S* values uncorrected for background, since in mpH^2^MM all photons within bursts are taken into

**Figure 4—figure supplement 3.**
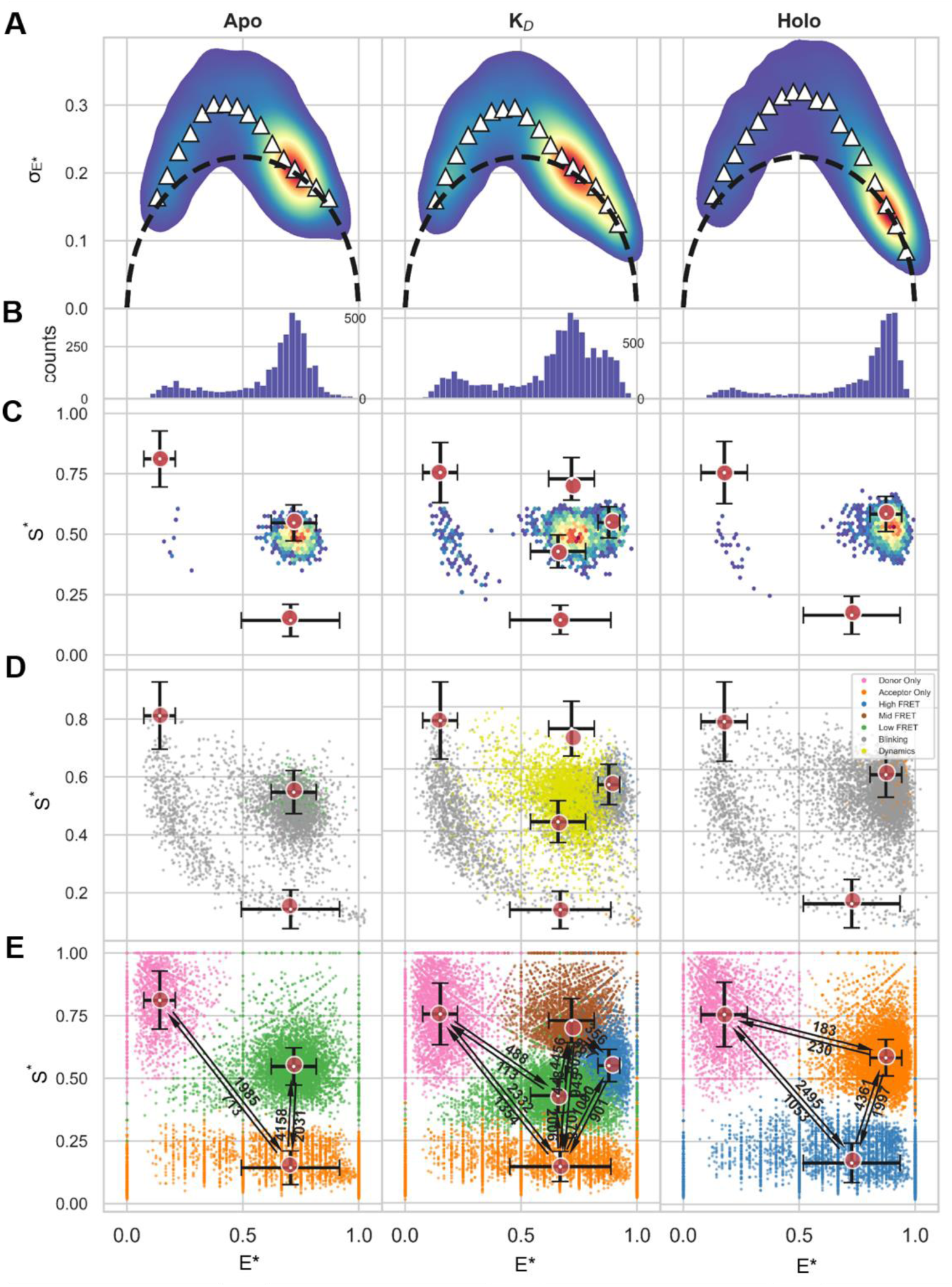
**Screening GlnBP(59C-130C) for rapid within-burst FRET dynamics**. Confocal-based single-molecule FRET results for GlnBP doubly-labeled at residues 59 and 130 with AF555 and AF647, in the apo state, near the K_D_, and holo state. (A) Burst variance analysis showing a weak signature of within-burst FRET dynamics. (B) Histograms of E* values of bursts, (C) E* versus S* 2D histograms of bursts, (D) 2D scatter plots of bursts classified by mpH^2^MM, with colors corresponding to which state(s) are present within the burst as determined with the Viterbi algorithm. Locations of states are given by red circles, and black crosses represent the SD of E* and S* values of dwells within each state. (E) E* versus S* 2D scatter plots of of dwells in mpH^2^MM-detected states within bursts detected by the Viterbi algorithm. Red circles and black crosses are same as in (D). Arrows and adjacent numbers indicate transition rates in s^-1^ units. Transitions with rates less than 100 s^-1^ are omitted, since such slow transitions are improbable to occur within single-molecule bursts with durations shorter than 10 ms and are most probably a mathematical outcome of the mpH^2^MM framework. The dispersion of the E* and S* values of dwells in mpH^2^MM-detected states are due to the short dwell times in these states, where the shorter the dwell time in a state is, the lower the number of photons it will include, and hence the larger the uncertainty will be in the calculation of E* and S* values of dwells. E* and S* are E* and S* values uncorrected for background, since in mpH^2^MM all photons within bursts are taken into account, including ones that might be due to background.

**Figure 6—figure supplement 1.**
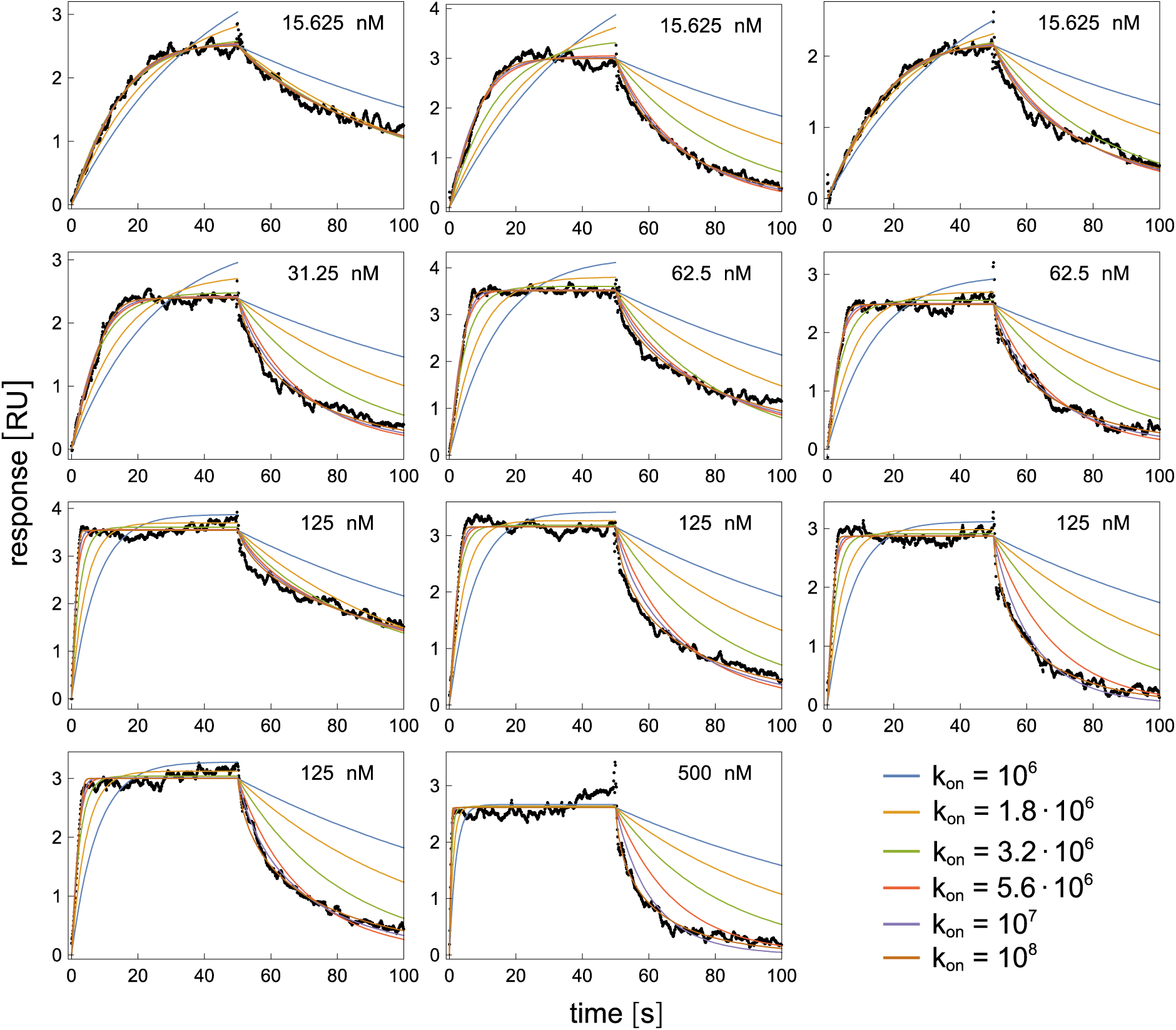
SPR sensorgrams at the indicated glutamine concentrations, and fits of the sensorgrams for different values of the effective on-rate constant, *k*_on_, as in Fig. 6B/C. The rescaled sum of squared residuals versus *k*_on_ for these fits is shown by dashed lines in Fig. 6D.

**Figure 7–figure supplement 1.**
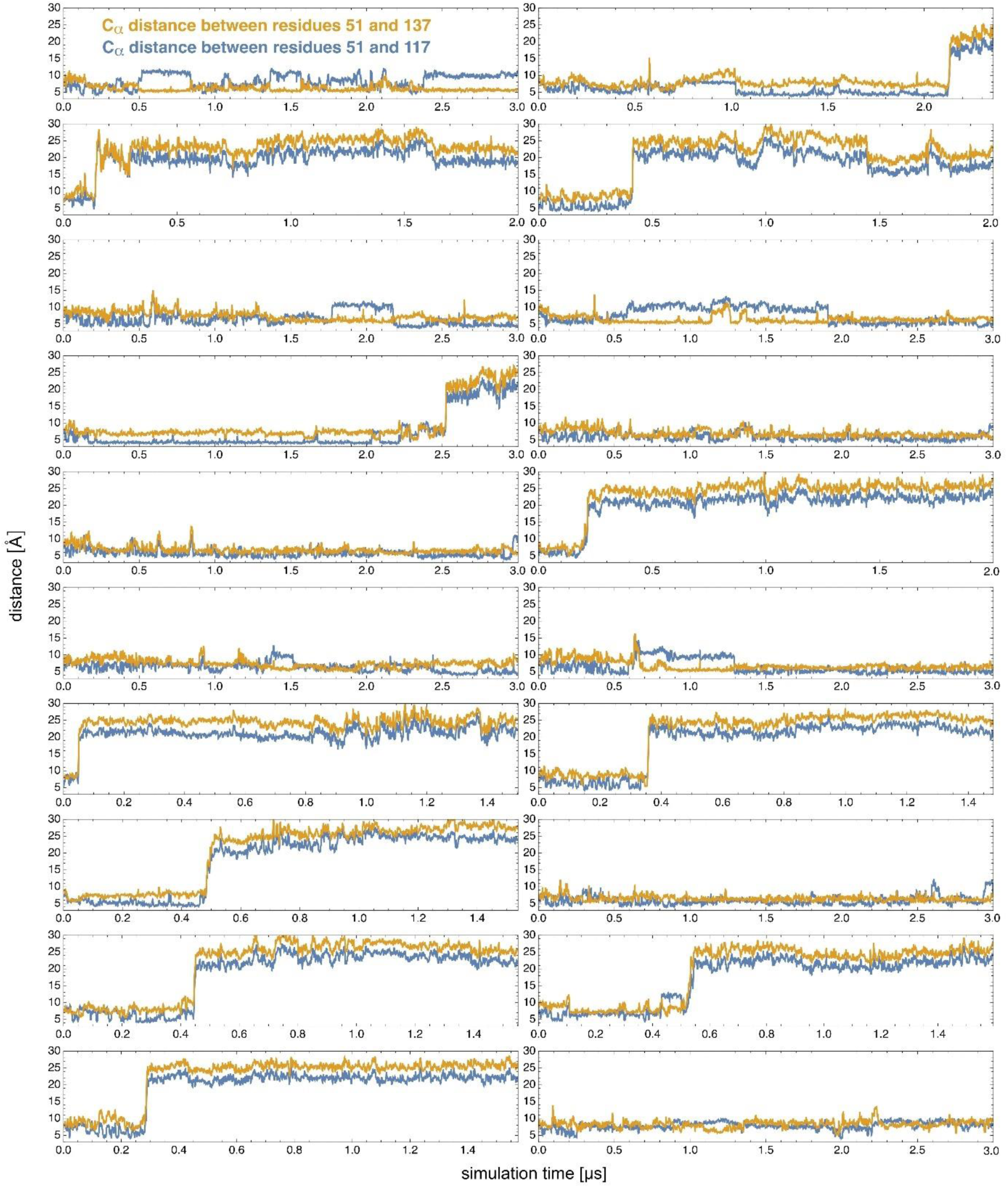
Opening transitions along MD trajectories starting from the closed, ligand-free protein conformation. Characteristic distances reflecting the protein conformation on 20 simulation trajectories with a length up to 3 µs starting from the closed but ligand-free conformation. On 11 of the 20 trajectories, rather sudden increases in the distance between the C_a_ atoms of the residues 51 and 117 (blue) and 51 and 137 (yellow) indicate a conformational transition from closed to open. On none of these trajectories, a transition back to the closed conformation occurs after opening. On the other 9 trajectories, GlnBP remains in the closed conformation for the entire trajectory length of 3 µs.

**Figure 8—figure supplement 1.**
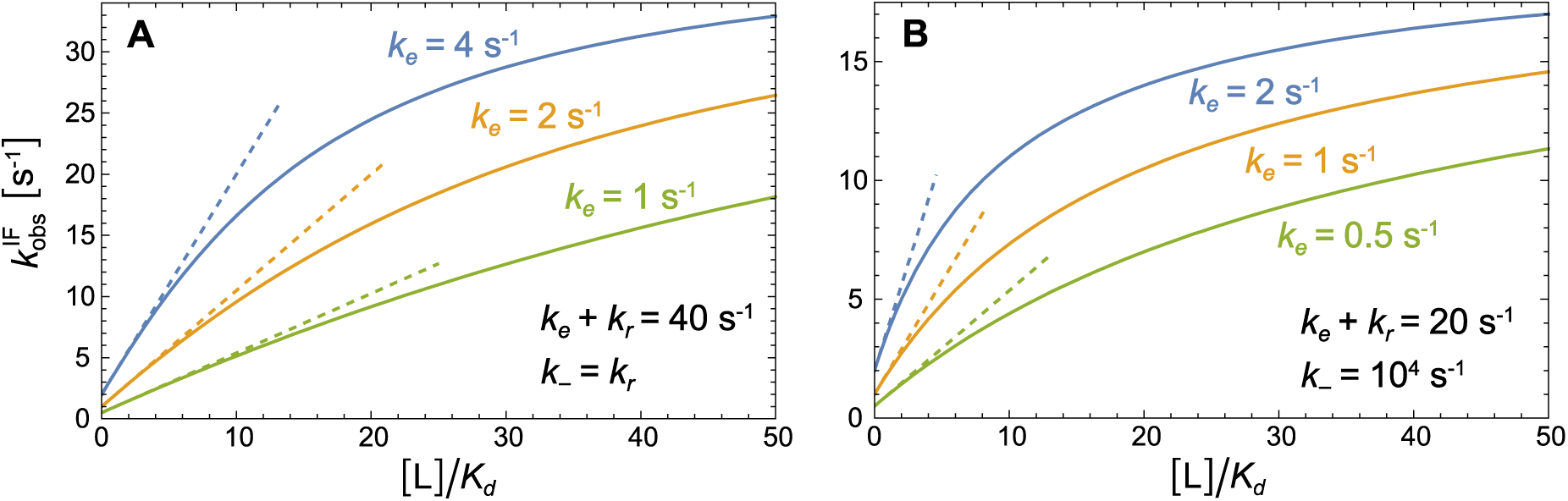
Exemplary plots of the dominant relaxation rate *k*^IF^ versus ligand concentration [L] for rate parameters consistent with Eq. (7). Full lines represent the exact solution given in the caption of Fig. 7, and dashed lines represent the approximate solution for sufficiently small [L] based on Eq. (4). In (A), the effective off-rate resulting from the exemplary parameters is *k*_off_ = 0.5 *k*_e_. In (B), the effective off-rate is *k*_off_ ≈ *k*_e_ because the unbinding process is dominated by the opening of the closed ligand-bound conformation with rate *k*_e_ for *k*_−_ ≫ 20 *k*_*r*_ as in this example. The limiting value of *k*^IF^ at large ligand concentrations is *k*_e_ + *k*_*r*_.

### Appendix 1: Interpretation of mpH^2^MM analysis

For analysis of within burst dynamics, we used multi-parameter photon-by-photon hidden Markov modelling (mpH^2^MM)^2, 3^ to identify the most-likely state model that describes the experimental results based on how E* and S* values may change within single-molecule bursts. For this analysis we (i) report the most-likely number of states and their mean E* and S* values (Fig. 4B, red dots). (ii) We investigate whether molecules traversing the confocal excitation volume are fully static and only in the mid-FRET state or high-FRET state, or whether they undergo dynamic FRET changes including transitions of mid/high-FRET states with photo-blinking dynamics or dark donor or acceptor states (Fig. 4B). (iii) We finally report on E* and S* values for parts of bursts with dwells in one of the identified states and the rate constants of transitioning between them (Fig. 4B). These analyses confirm that among the two types of dynamic transitions that influence the burst-based E* and S* values, these are mostly donor or acceptor photo-blinking dynamics between bright and dark states of the fluorophores. Such behavior is irrelevant to understanding the conformational changes in GlnBP but does influence the mean FRET efficiency values if not decoupled. Importantly, no dynamic transitions occur between the mid-FRET and high-FRET states at timescales shorter than 10 ms (i.e., with rate constants higher than 100 s^-1^). All measurement conditions show significant photo-blinking dynamics which occur mostly on few ms to sub-millisecond timescales most prominently for the use of AF555/AF647 and the GlnBP(59/130) variant (compare Fig. 4 and Figure 4—figure supplement 1-3). Therefore, the blinking dynamics likely account also for the signature of within-burst dynamics shown by BVA (Fig. 4, Figure 4—figure supplement 1-3).

Most importantly, mpH^2^MM identifies single apo and holo E*-states, which describe the open mid-FRET and closed high-FRET conformations of GlnBP. Only in the presence of low (near K_D_) concentrations of glutamine two FRET states are identified which interconvert on timescales slower than 10 ms. Notably, the mean E* and S* values of the FRET states are slightly dissimilar to the centers of the burst-based E* and S* populations, owing to the effect of the rapid photo-blinking dynamics within bursts, which lead to averaging the E* and S* values of the FRET states with those of the photo-blinked states. Additionally, in the presence of near-K_D_ concentrations of glutamine, the FRET dynamics occur in the few ms timescale or even slower, which may contribute only slightly to the signature of FRET dynamics in BVA. In conclusion, if intrinsic conformational dynamics existed in apo GlnBP, it could only be between the highly-populated FRET conformation we identify and another conformation that is populated way below the sensitivity of our measurement and analysis (potentially <5-10% populations). Thus, we can conclude that the majority of the conformational dynamics in GlnBP is induced by glutamine, most probably as a result of its binding to GlnBP.

### Appendix 2: Description of TIRF data acquisition and analysis

At first, we studied a biotin-modified double-stranded DNA (dsDNA), which was labeled with Cy3B (donor) and ATTO 647N (acceptor) in 13 bp distance, and used this as a reference sample to allow a direct comparison of µsALEX and TIRF data (Appendix Figure 2). For this, we immobilized the dsDNA on a PEG-coated glass surface via streptavidin-biotin interactions. We recorded both donor and acceptor fluorescence via a dual-view split on our EMCCD camera with 100 ms integration time per frame. With this we obtained traces that lasted multiple 10 second periods. Since we did not perform millisecond alternation of green-and-red laser excitation, we verified that the sum-signal of the donor and acceptor channel was constant as a function of time for each molecule and discarded traces that did not obey this condition. The dsDNA sample displays an apparent FRET efficiency E* of ∼0.64 for in-solution measurements, which agreed well with the analysis of surface-immobilized molecules on the TIRF microscope having a mean E* of 0.62 (Appendix Figure 2A/B).

Then, we investigated the conformational states and changes of GlnBP(111C-192C) with the dye pair ATTO 532/ATTO 643, since these showed least photophysical FRET-dynamics (see main text and Appendix 1). To exclude the influence of buffer and other small molecules in TIRF measurements on the conformational state of GlnBP, we initially performed control experiments in µsALEX (Appendix Figure1). We found that GlnBP was influenced by the addition of oxygen scavenger cocktails (pyranose oxidase and catalase, POC, and glucose or protocatechuate-dioxygenase, PCD, and 3,4-protocatechuicacid, PCA), resulting in the formation of artificial holo-state GlnBP molecules (Appendix Figure 1E/F). In TIRF experiments, the effect of oxygen scavenger might have been misinterpreted as intrinsic closing. We consequently proceeded with no oxygen-removal in PBS buffer (pH 7.4) and with 2 mM Trolox as photostabilizer. GlnBP was immobilized by biotin-NTA interactions mediated by Nickel(II). To our surprise we found very different mean E* values on TIRF in comparison to μsALEX measurements (Appendix Figure 2E/F). In detail, the mean E* values were much higher on TIRF than on µsALEX (Appendix Figure 2E/F) in contrast to dsDNA (Appendix Figure 2A/B). This can be interpreted as an altered conformational state of GlnBP, e.g., likely caused by protein-glass interactions due to surface-immobilization or interaction of the protein or dyes with the biotin-NTA moiety. Furthermore, addition of saturating glutamine concentrations did not show the expected behavior of a full shift of the population to a higher-FRET state (Appendix Figure2F). Instead, only a small fraction of the population is shifted for both low and saturating glutamine concentrations. At concentrations of glutamine around the K_d_-value freely-diffusing GlnBP shows a mix of open- and closed state in µsALEX experiments (Fig. 3). In TIRF, however, we could not identify dynamic transitions (Appendix Figure 2/4). This finding indicates that a part of the immobilized fluorophore-labelled GlnBP becomes non-functional. Since our protocol deviates from that used in other studies^4–6^, we probed whether we could reproduce published data on substrate-binding domain 1 and 2 (SBD1 and SBD2)^7^. Again, we find a good match between biochemical properties, µsALEX and the corresponding TIRF data for both proteins (Appendix Figure 5).

**Appendix figure 1.**
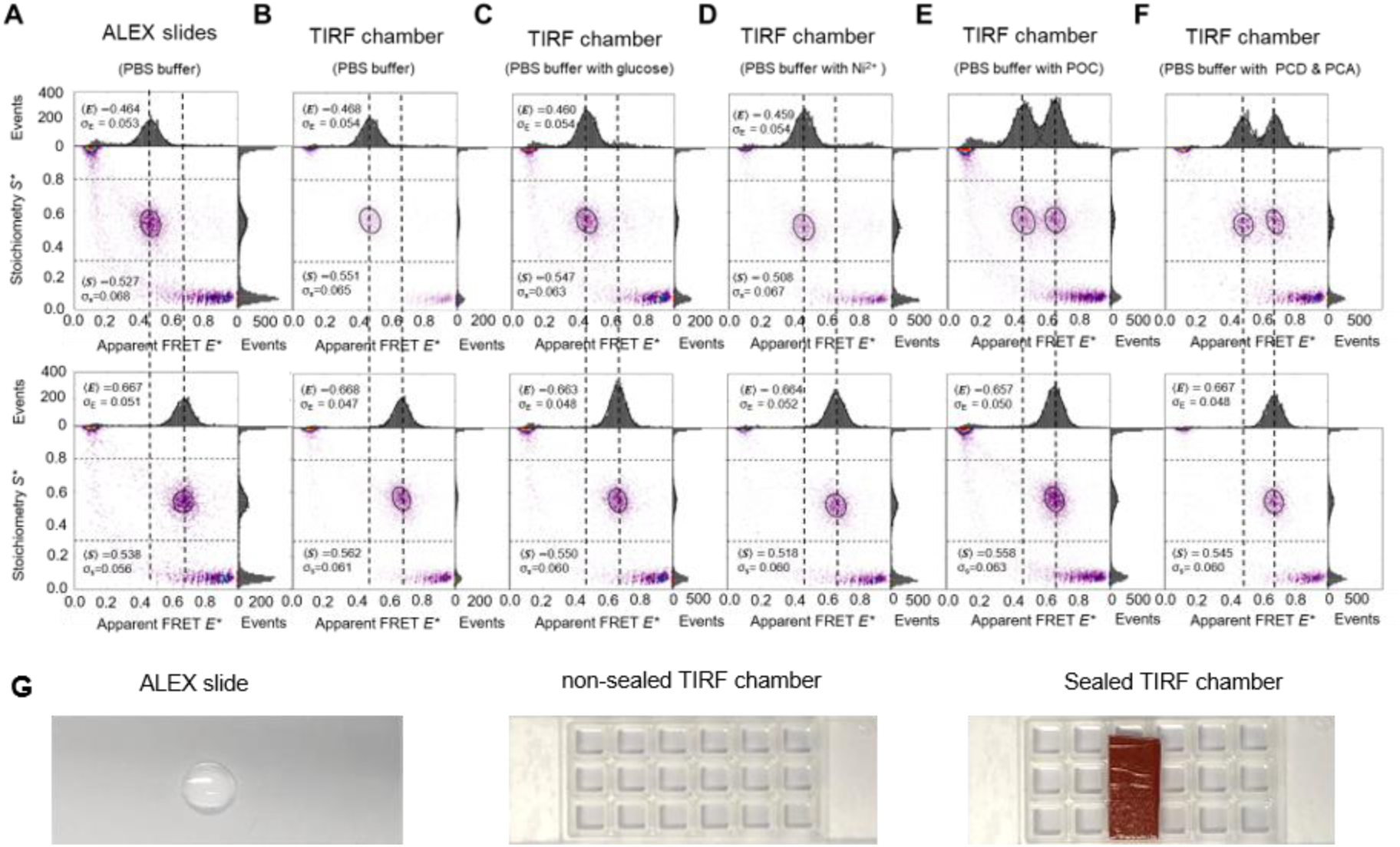
Buffer effects on the conformational states of GlnBP(111C-192C) under various conditions. Due to the high binding affinity of GlnBP for L-glutamine, several control experiments under different conditions were performed to exclude artifacts induced by the reagents present in each set of experiments. The µsALEX experiments of the refolded GlnBP(111C-192C) double-cysteine variant labeled with LD555/LD655 fluorophore pairs were measured in PBS buffer (pH 7.4) using conventional microscope glass slides (A) and using TIRF chamber (B). The PBS buffer containing (C) 40 mM glucose, (D) 50 nM Ni^2+^, (E) pyranose oxidase/catalase (POC) and (F) protocatechuate-dioxygenase (PCD)/3,4-protocatechuicacid (PCA) was used for the ALEX measurements. (G) The conventional glass coverslips used in µsALEX experiments (left figure) and TIRF chambers (sticky-Slide 18 well, Ibidi; non-sealed chambers: middle panel; sealed: right panel) glued on top of PEG-/biotin-PEG-silane microscope glass coverslips used in the TIRF experiments.

**Appendix figure 2.**
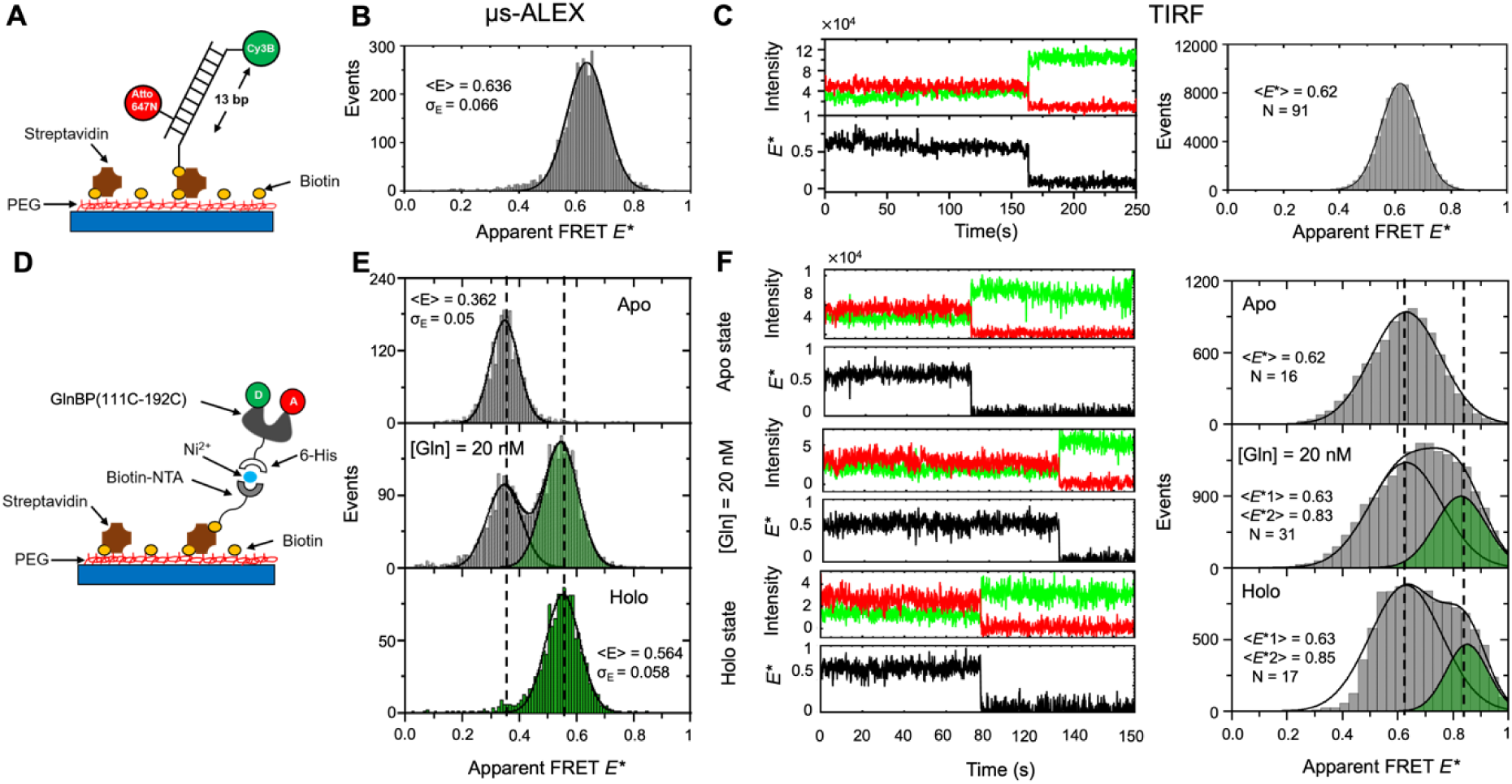
Comparing smFRET measurements of biotin-modified dsDNA and GlnBP(111C-192C) using diffusion-based µsALEX versus TIRF microscopy. (A) Schematic view of dsDNA labeled with Cy3B and ATTO 647N for smFRET characterization on PEGylated coverslips. (B) Typical μsALEX-based E*-S* histograms of the biotin-modified dsDNA labeled with Cy3B and ATTO 647N. (C) Representative fluorescence time trace of respective single emitter of the biotin-modified dsDNA sample under continuous wave excitation of ∼500 µW at 532 nm and the FRET histograms of all analyzed molecules and the FRET histograms of all measured molecules combined. (D) Schematic view of the refolded GlnBP(111C-192C) labeled with ATTO 532 and ATTO 643 for smFRET characterization. (E) Typical μsALEX-based E*-S* histograms of the refolded GlnBP(111C-192C). (F) Representative fluorescence time trace of respective single emitter of the refolded GlnBP(111C-192C) under continuous wave excitation of ∼500 µW at 532 nm and the FRET histograms of all analyzed molecules. Additional data for each condition is shown in Fig. S13/S14.

**Appendix figure 3.**
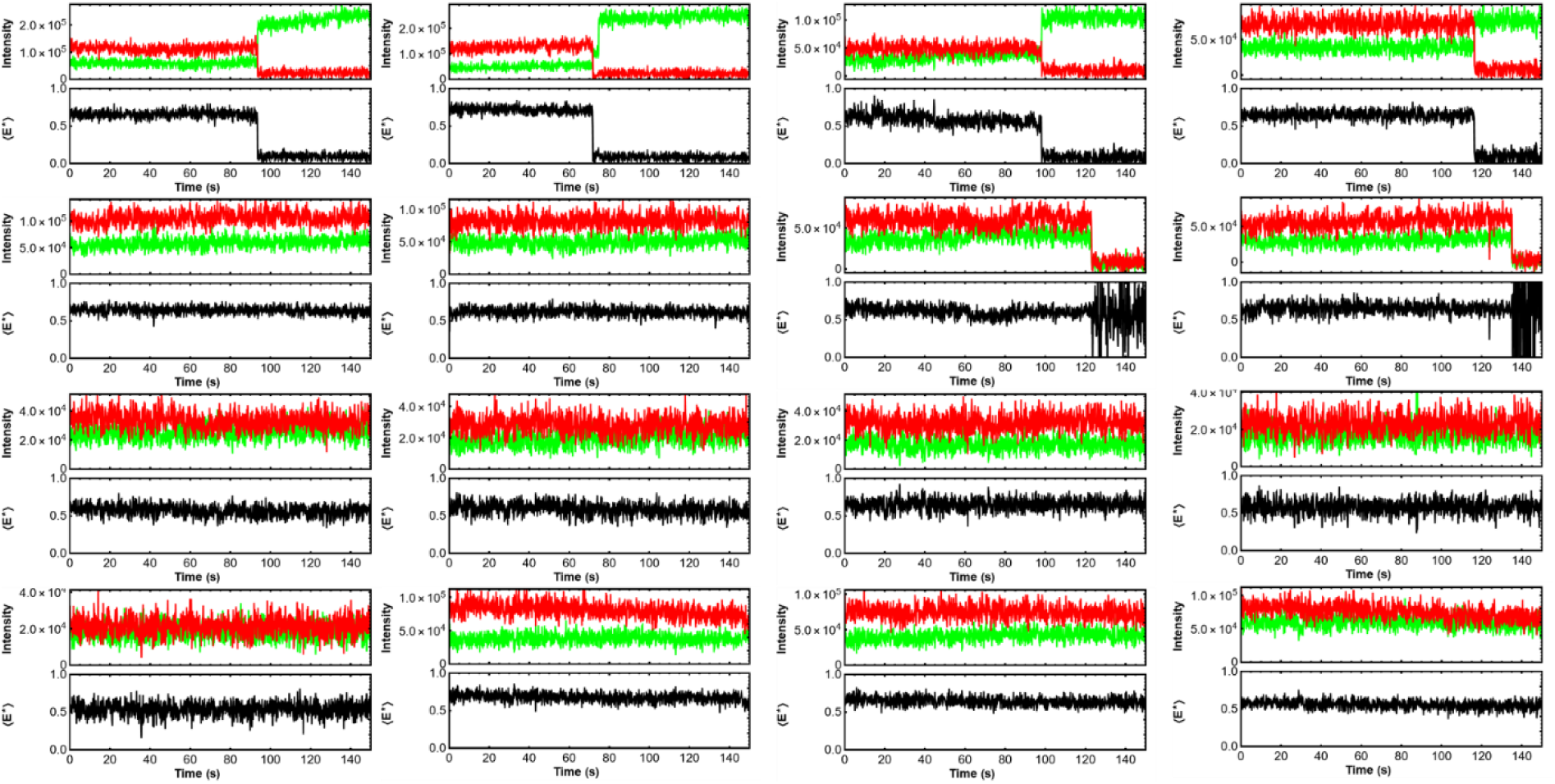
Representative fluorescence time traces of respective single emitter of biotin-functionalized DNA labeled by maleimide-modified derivatives Cy3B and ATTO 647N (13 bp inter-dye distance). All measurements were done in oxygen scavenging buffer (3 U/mL of pyranose oxidase, 90 U/mL of catalase and 40 mM glucose, PBS buffer, pH 7.4). Laser power: 500 µW.

**Appendix figure 4.**
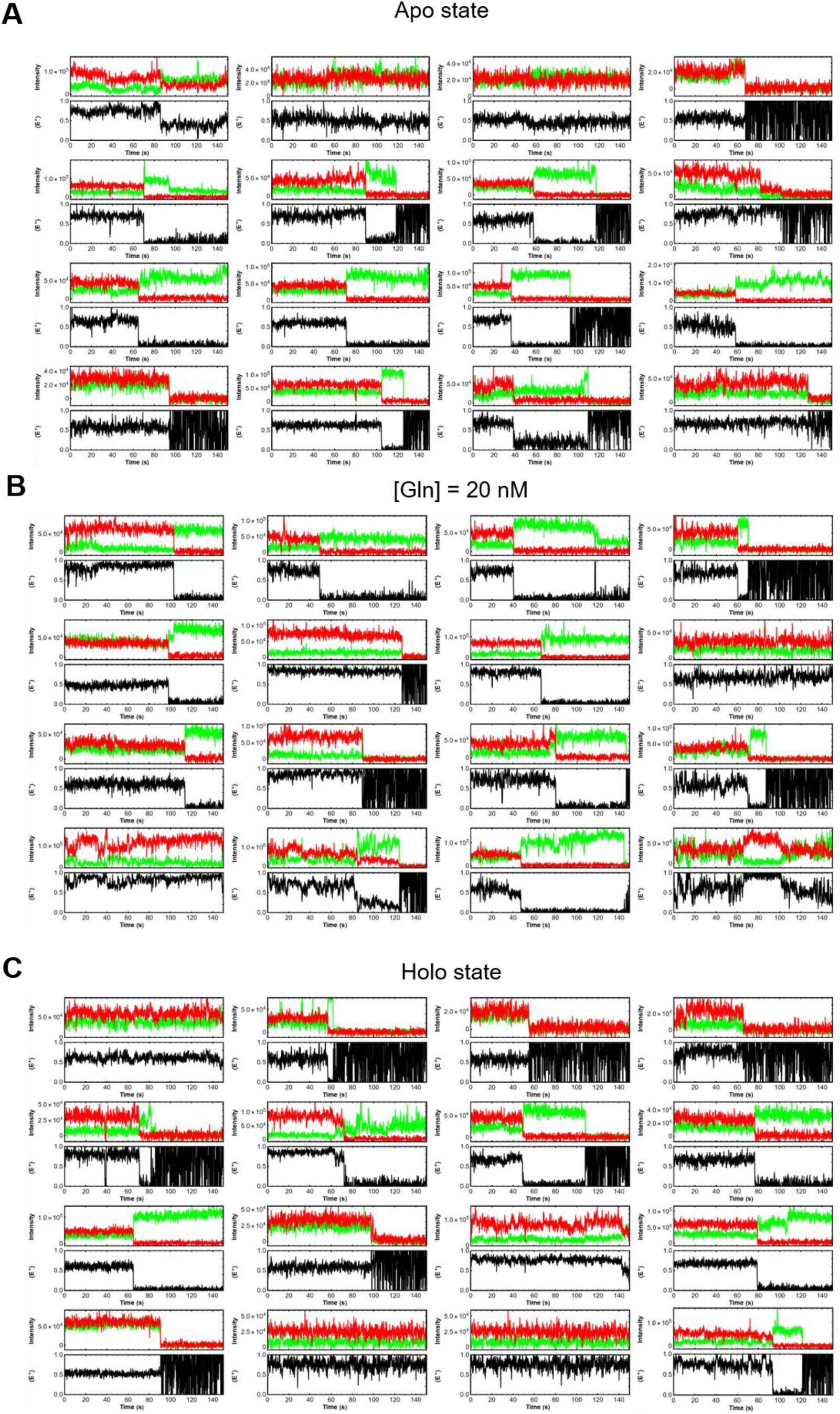
Examples of fluorescence time traces of respective single emitter of refolded GlnBP(111C-192C) labeled by maleimide-modified derivatives ATTO 532 and ATTO 643. All measurements were done in PBS buffer, pH 7.4 and 2 mM Trolox. Laser power with continuous 532 nm excitation: 200 μW.

**Appendix figure 5.**
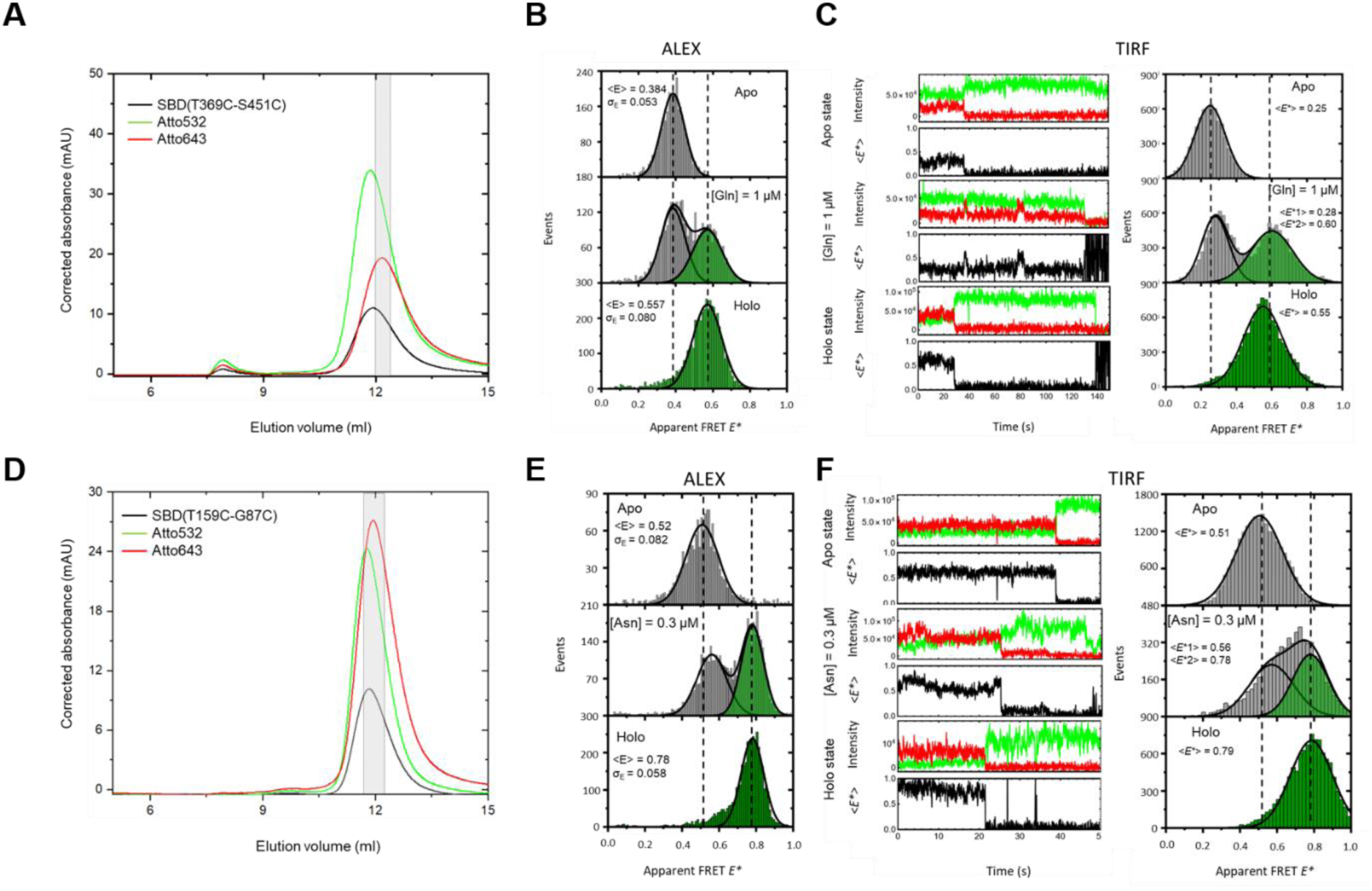
(A) and (D) Size Exclusion Chromatography (SEC) of SBD(T369C-S451C) and SBD(T159C-G87C). The selected fractions (grey-shaded area) were collected and used for the solution-based smFRET measurements. The selected fractions (grey-shaded area) having the best overlap of protein, donor, and acceptor absorption were used. The protein absorption was measured at 280 nm (black curves) and the donor dye (ATTO 532) absorption at 532 nm. The acceptor dye absorption (red lines) was measured at 643 nm for ATTO 643. (B) and (E) Typical μsALEX-based E*-S* histograms of the SBD(T369C-S451C) and SBD(T159C-G87C). (C) and (F) Representative fluorescence time trace of respective single emitter of the SBD(T369C-S451C) and SBD(T159C-G87C) and the FRET histograms of all measured molecules.

### Appendix 3: Considerations on the accessibility of the ligand binding pocket for solvent and ligand in the closed conformation of GlnBP

To describe the expected binding behavior of the substrate glutamine to GlnBP, we performed docking calculations of GlnBP in its open and closed conformations. The GlnBP structure that represents the open conformation is the one reported under pdb code 1GGG^10^. The GlnBP structure that represents the closed conformation is the one reported under pdb code 1WDN^1^, with the bound ligand taken out of the file. Then, we used the 3D conformer structure of the ligand to be docked onto the structures of GlnBP. We used the SwissDock web server to perform the docking procedure^11, 12^. The results show that (i) while glutamine can dock to many sites on GlnBP, the results that yield the lowest binding free energy are when it docks onto its cognate binding site, both in the open and closed conformation (Appendix Figure 1-2). (ii) The calculated binding free energy of Gln to GlnBP in the optimized docking site leads to a dissociation constant of 20 µM in the open conformation and 230 nM in the closed conformation (Appendix Figure 2), about two orders of magnitude different. (iii) The higher binding free energy is due to the larger amount of GlnBP residues when the docked glutamine interacts with in the closed conformation relative to in the open conformation. (iv) The binding pocket in GlnBP seems to surround the docked glutamine from all directions (Appendix Figure 1), which implies that it is less probable that glutamine can access the binding pocket in the closed conformation. Instead, it is more probable that the glutamine reaches its binding site in GlnBP when it is not yet closed.

**Appendix figure 1.**
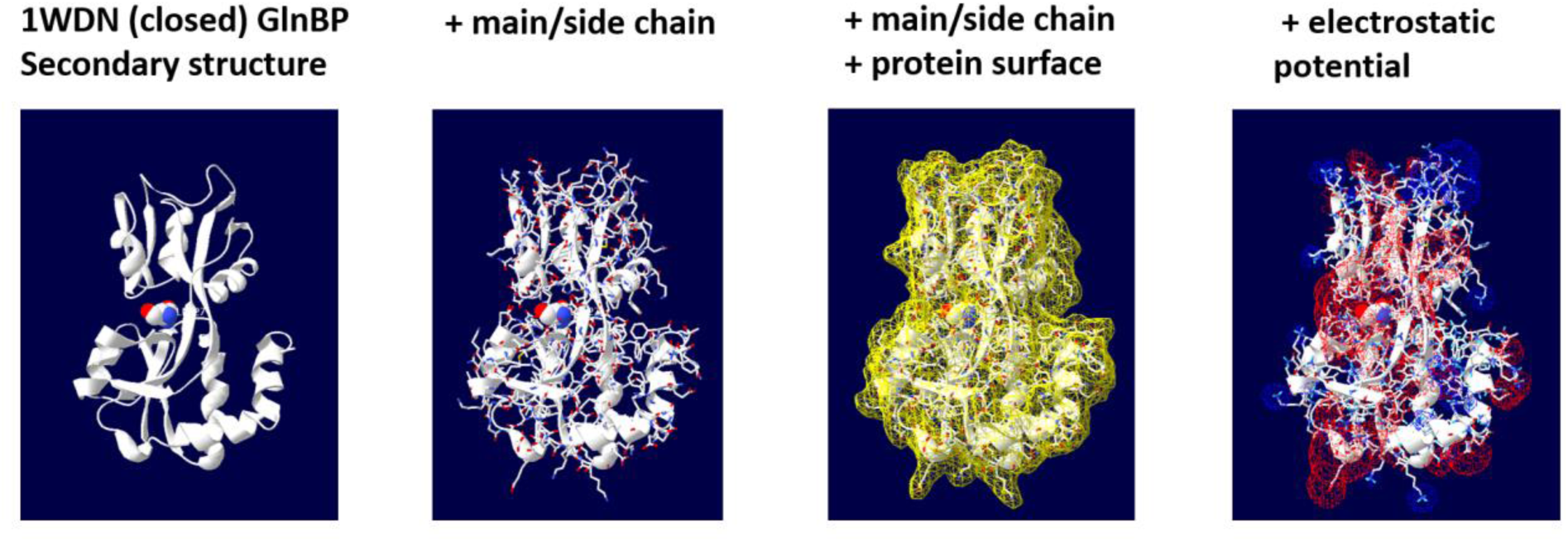
The structure of holo GlnBP with optimized docking of glutamine. The figure reports the optimized results of docking glutamine onto the crystal structure of GlnBP in holo form, after the glutamine substrate was removed from the structure, and presented back as a docking ligand using the SwissDock web server. From left to right: (i) the glutamine is docked onto the correct binding pocket within the closed conformation of GlnBP, (ii) amino acid side chains are wrapping the docked glutamine from all directions, (iii) and indeed the protein surface covers the docked glutamine, and (iv) the residues covering the docked glutamine seem to carry a net negative charge.

**Appendix figure 2.**
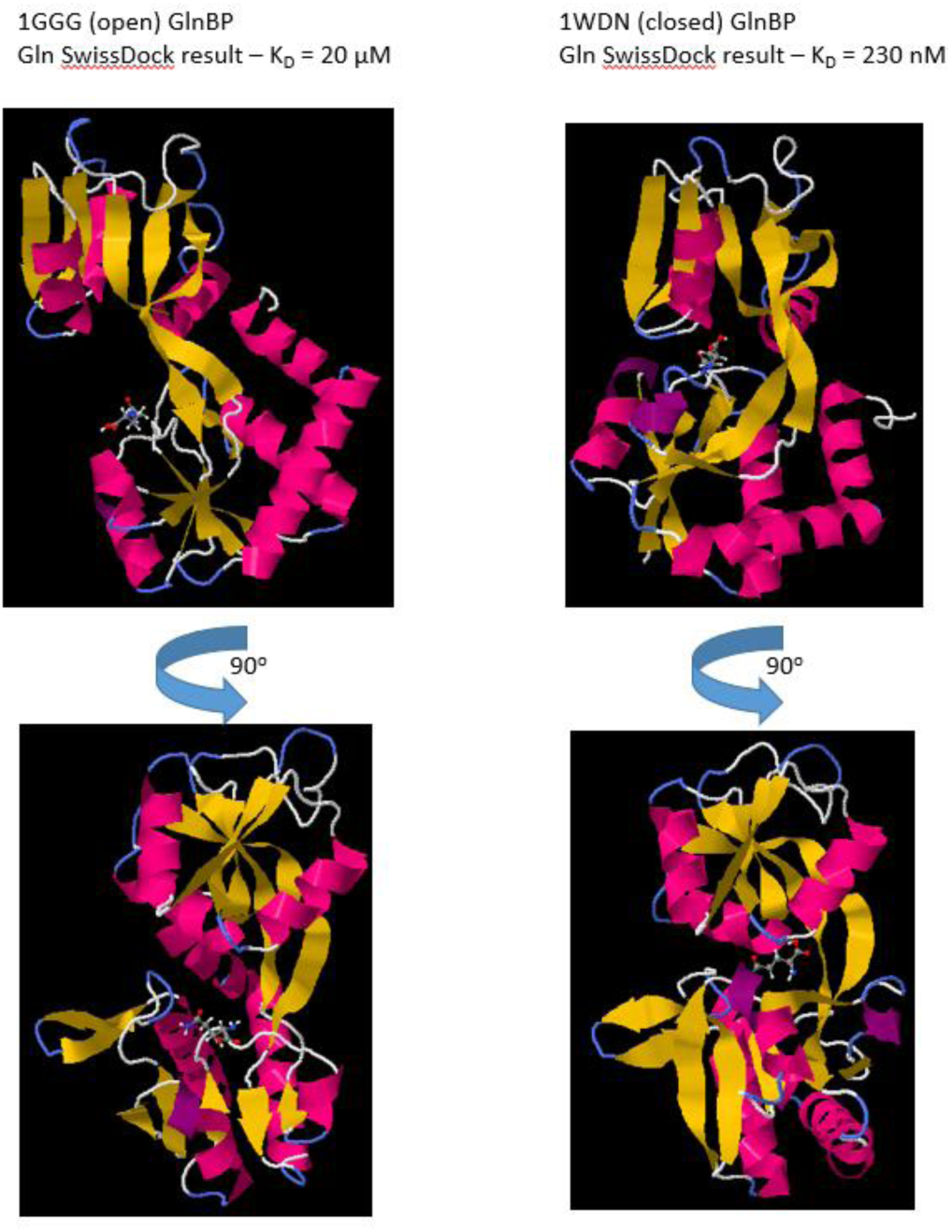
Optimized docking of glutamine to GlnBP in its open and closed conformations. Using the SwissDock web server, the molecule glutamine was docked onto the crystal structures of GlnBP in its open (pdb:1GGG) and closed (pdb:1WDN; with the glutamine substrate taken away) conformations, and the optimized docking sites as well as the calculated dissociation constant are shown (dissociation constant is calculated out of the binding energies reported in the docking results). The preferred docking of glutamine is the same site within GlnBP. The difference is that while in the open conformation glutamine binds to one domain with the other as a distant domain, in the closed conformation the other domain closes on top of the docked glutamine . Following the calculated binding energies from the optimized docking results, while the dissociation constant of glutamine to GlnBP is 20 µM in the open conformation, in the closed conformation it is 230 nM.

